# Methane-producing microorganisms are widespread in surface waters and floating algal mats of inshore Baltic Sea habitats

**DOI:** 10.64898/2026.07.27.740719

**Authors:** Maysoon Lundevall Zara

## Abstract

Inshore coastal waters are almost invariably supersaturated with respect to methane and are thereby sources of methane to the atmosphere. We investigated floating algal mats and surface waters of four contrasting inshore habitats and quantified methane concentrations, sea-to-air emissions, and microbial community composition of surface waters over a seasonal cycle to determine the potential for in-situ microbial methane production in shallow oxygen-saturated surface waters with floating algal biomass.

16S rDNA sequencing indicated that Archaea belonging to the genera Methanocorpusculum, Methanosarcina, Candidatus Methanomethylophilus, and some genera from order Methanobacteriales occurred in the floating algal mats. qPCR of the genes encoding the methyl coenzyme M reductase mcrA revealed the highest expression levels during the warmest sampling periods supporting active methane production directly in surface water. Co-occurrence of the Archaea sequences and sequences belonging to the cyanobacterium strain Nodularia PCC 9350 suggests a structural relationship. Our study underscores the significant, yet underexplored impact of methane production on the surface in aggregates of floating algal material. While Nodularia and methanogens can exist independently in surface waters, their co-occurrence in algal mats reveals where the layered mat structure creates distinct microenvironments that facilitate direct metabolic exchange and provide physical stability for both groups, thereby potentially enhancing methane production in these shallow coastal systems.

## Introduction

Oversaturation of dissolved methane in the marine surface mixed layer has been reported for a number of marine environments over the past two decades (Bižić et al., 2020b; Lundevall-Zara et al., 2021; Roth et al., 2023; Taenzer et al., 2020; Watanabe et al., 1995). The surface water concentrations are often higher than in corresponding deeper water below the thermocline and suggest a methane source that is produced either in-situ in the euphotic zone or transported laterally from a nearby methane-rich environment (Bižić et al., 2020b; Stawiarski et al., 2019).

The responsible microorganisms, metabolic pathways, and a causative link between photosynthesis and methane production remain unclear, despite several possible lines of evidence for methanogenetic pathways in oxic waters. Traditionally, microbial methanogenesis is an obligate anaerobic pathway, carried out by Archaea via three main pathways: hydrogenotrophic, methylotrophic, and acetoclastic (Conrad, 2020). The thermodynamic and kinetic constraints of hydrogenotrophic and acetoclastic methanogenesis make these pathways less likely for oxic methane production, although hydrogen transfer from photoautotrophic cyanobacteria to methanogens has been suggested (Grossart et al., 2011). Instead, there is growing acceptance for methylotrophy under oxic conditions and further non-competitive pathways of methanogenesis independent of the syntrophic consumption of fermentation products hydrogen and acetate such as the microbial degradation of C-P compounds and demethylation of methylphosphonate (Karl et al., 2008). For example, degradation of methylamines and methylsulfides (Damm et al., 2008) have been proposed as more likely candidates for microbial methanogenesis under oxic conditions (Taenzer et al., 2020; Ye et al., 2020).

Surface waters of inshore habitats with dense floating algal mats and benthic vegetation as sites of important methanogenesis have not been considered in previous studies. Protected coastal embayment of the Baltic Sea often develop thick mats of floating macroalgal mats dominated by the chlorophyte Cladophora glomerata during the summer (Lehvo and Bäck, 2001). These surface waters in these habitats exhibit high methane concentrations up to 250 nmol/L and are strong sources of methane to the atmosphere, particularly during the warmest period of the year (Bižić et al., 2020b; Björk et al., 2023; “Coastal ecosystems, reservoirs of life | ICOS,” n.d.). Floating macroalgal biomass are common in eutrophic, productive inshore waters and represent close associations of autotrophic and heterotrophic organisms (Lyons et al., 2014; Global Change Biology) or as an alternative metabolic process involving methylphosphonate or other methylotrophy, or within anoxic niches in dense, decaying algal matter, and these habitats are prevalent in inshore waters (Bange et al., 1994; Izhitskaya et al., 2022; Weber et al., 2019). Similar to macrophyte beach wrack on Baltic seashores (Björk et al., 2023), floating macroalgal aggregates may be significant sites for surface water methanogenesis.

Submerged macrophytes can serve as hosts for methanogenic archaea within their tissues, enabling methane production in anaerobic microsites inside the plants. These macrophytes can also create habitats that support various aquatic organisms capable of methane production through classic methanogenesis, even in areas where oxygen is present (Hilt et al., 2022).

The objective of this study was to determine the presence and activity of methanogenic Archaea in floating algal mats and surface waters of shallow inshore Baltic Sea habitats, and to assess whether these surface environments represent an important niche for methanogens and methane production. We aimed to establish a taxonomic classification of microbial communities in these ecosystems by analyzing both eukaryotes and prokaryotes through amplicon sequencing of particle size-fractionated water samples and macroalgal mat samples from contrasting habitats around Askö Island in the Trosa archipelago. Specifically, we sought to identify methanogens and determine the relative abundance patterns of these taxa using the functional biomarker gene mcrA (methyl-coenzyme M reductase) (Thauer et al., 2008), which is universally present in methanogenic Euryarchaeota. By combining taxonomic data with qPCR analysis of mcrA gene expression and correlating these findings with methane measurements, we investigated the relationship between methanogenic gene expression levels and the abundance of methane-producing microorganisms, ultimately seeking to understand methane sources and methanogenic pathways in shallow inshore habitats where algal mats create anoxic microenvironments favorable for archaeal growth.

Eukaryotes and prokaryotes were analyzed taxonomically by amplicon sequencing in particle size-fractionated water samples and macroalgal mat samples from contrasting inshore shallow-water habitats of the central Baltic Sea. In combination with qPCR analysis of the mcrA gene expression (an indicator of methane production), which combined with the taxonomic data helps understand methane sources and methanogenic pathways in shallow inshore habitats.

## Materials and methods

### General approach and sampling region

The southwestern end of the island is directed toward the open Baltic Sea and consists mainly of shallow embayment and rocky cliffs. The north-eastern end faces the calmer Trosa archipelago. It is a good example of a typical Baltic Sea inshore habitat.

The samples for this study are from inshore waters on the island of Askö in the Baltic Sea (Lundevall-Zara et al., 2021). They were chosen to represent four types of habitats based on a categorization described in (Lundevall-Zara et al., 2021). The categorization of the habitats is based on the water depth, amount of wind exposure, sediment type, human perturbation due to reed cutting, indicator species for eutrophication, occurrence of hydrogen sulfide, presence of floating algal mats, and occurrence of rising methane bubbles. The four different habitats A-D have been shown to be distinct with respect to methane emissions (Lundevall et al., 2021). Eight sampling stations have been selected from the 24 sampling stations of an earlier study from 2019 (Lundevall-Zara et al., 2021).

The characterization of Habitat A encompasses sheltered bights with water depths of up to 0.5 meters. These bights are characterized by reeds and soft, organic-rich clay, with a shoreline populated by deciduous trees. Along the shoreline, there are dense reed clusters, and during the months of July and August, substantial thick layers of algal mats are present. Human activities, such as agriculture and reed cutting, have also had an impact on this habitat. Notably, in June and August, the habitat experiences the emergence of bubbles accompanied by the distinctive odor of hydrogen sulfide.

Habitat B shares similarities with Habitat A, featuring soft, organically rich sediment. However, it differs in terms of water depth, ranging from 0.5 to 1.0 meters, and comprises open bights. The shoreline is predominantly vegetated with deciduous trees, and this habitat remains unaffected by human activities. Approximately half of the area is covered by benthic algae, and no visible bubble emissions occur in this habitat.

Habitat C is characterized by sandy sediment and unvegetated rocky terrain, with water depths of up to 2.0 meters. It is situated in more open areas, subject to higher wave activity and currents. Notably, this habitat lacks algal mats and sessile aquatic vegetation, and there is no significant presence of hydrogen sulfide odor.

Habitat D consists of hard, unvegetated ground and strong currents, with water depths ranging from 0.5 to 1.5 meters. Its shoreline features rocky cliffs with sparse vegetation, including sessile aquatic plants. During the summer months, algal mats accumulate in this habitat. Importantly, there is no discernible odor of hydrogen sulfide, nor are visible bubbles present (Figure 1).

**Figure 1.**
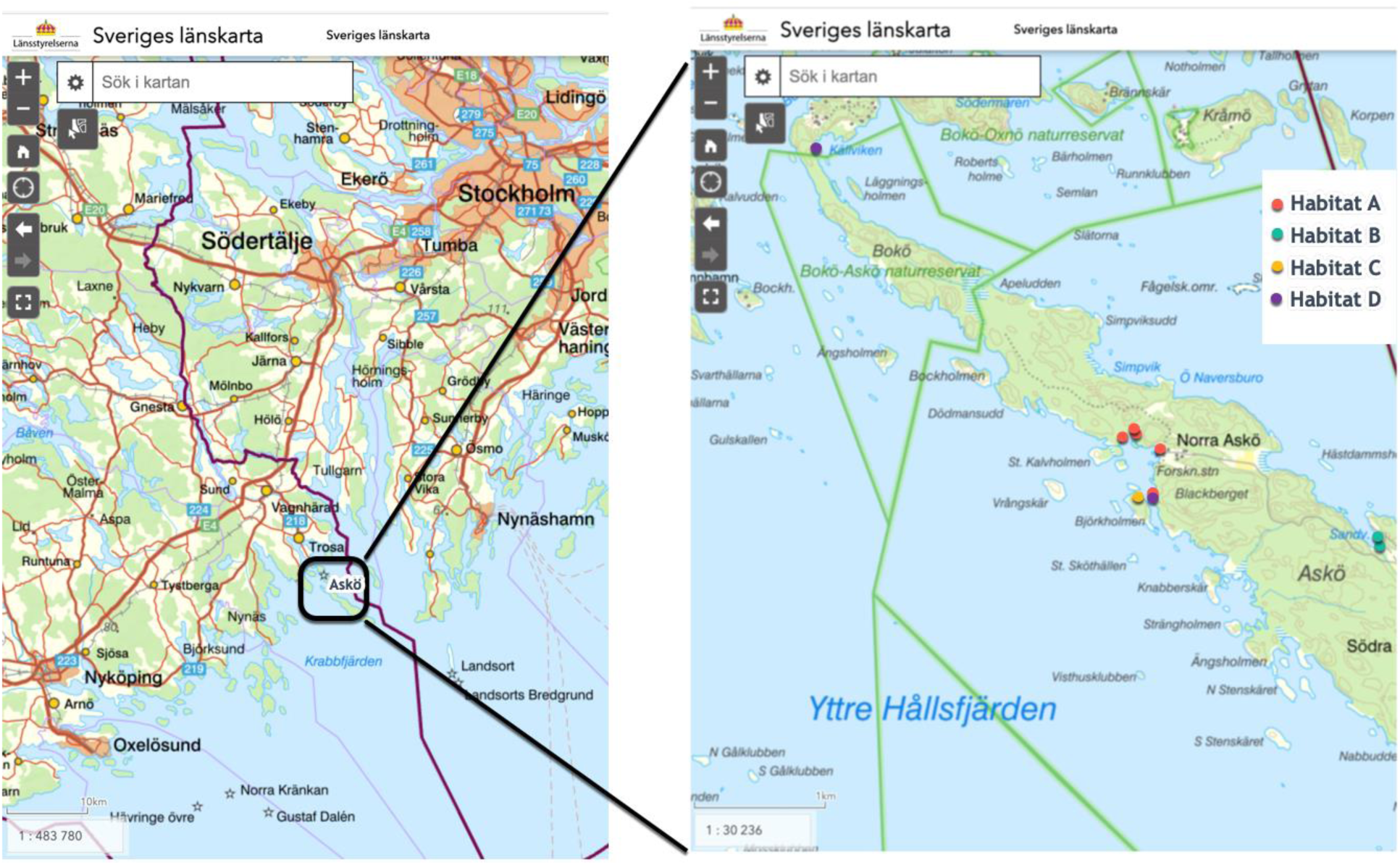
Map of sampling locations in Askö Island grouped by habitats.

### Field Methods

On each sampling occasion and before collecting each sample, the following data were recorded: air and water concentration of methane, date, time, salinity in water, air and water temperature, O_2_ concentration in water, and GPS location. Wind velocity data and air temperature were obtained from the database of the Swedish Meteorological and Hydrological Institute (“Askö Kustmätsystem, SMHI,” n.d.). The concentration of dissolved O_2_, water temperature, and salinity data were obtained with a handheld WTW meter.

Water samples were collected in a 5-liter bucket, from the eight chosen stations to filter later in the lab, while algal mat samples were collected directly in plastic bags for samples and kept in dry ice in an ice box until they were placed in –80°C freezer in the lab.

There were 19 algal mat samples collected from habitats A, B, and D. Samples were collected directed from the water surface and stored in plastic sample bags in flash-frozen in liquid N_2_ and stored at –80°C until extraction of DNA and RNA.

Methane concentrations were determined using water samples collected in a 120 ml glass vial filled and crimp closed with a butyl rubber stopper and aluminum caps without gas bubbles. For each 1ml volume water, 0.1 ml Zinc chloride, or 1% ZnCl_2_ of total volume had been added. The glass vials were kept upside-down until all vials were analyzed with Gas Chromatograph (GC) in the lab.

### Laboratory methods

#### Methane Analysis and calculation of methane flux

Dissolved methane concentrations were measured using headspace equilibration followed by gas chromatography with FID detection. Methane fluxes were calculated using two approaches: (1) floating chamber measurements with 8L chambers deployed for 15-78 hours, both with and without bubble shields to quantify ebullitive versus diffusive fluxes, and (2) a boundary layer gas transfer model incorporating dissolved methane concentrations, wind velocity, water temperature, and salinity. The gas transfer velocity was calculated using the Wanninkhof, (2014) relationship, and chamber fluxes were determined from the linear increase in headspace methane concentration over time, as described in (Lundevall-Zara et al., 2021).

### Sample filtration

Sequential water filtration was performed with a peristaltic water pump Masterflex® Ismatec® Reglo Independent Channel Control (ICC) Peristaltic Pumps, Avantor® at a pump rate of 70ml/min using 25 mm Swinnex filter holders preloaded with PALL Supor membrane filters of pore sizes 10µm and 0.22µm to obtain two different size sample size fractions. 1–2 liters of water were filtered through the 10µm filter followed by filtration of 300 ml through the 0.22µm filter. The 10µm fraction separated zooplankton, colonial eukaryotes, and particle-associated prokaryotes from the free-living (0.2µm) fraction. We used 4-6 filter samples to obtain replicates for DNA and RNA extraction. The filters were transferred directly into Thermo Scientific Nunc CryoTubes, flash-frozen in liquid N_2_, and stored at –80°C until extraction of DNA and RNA.

### DNA and RNA extraction and sequencing

DNA was extracted using DNeasy Power Water kit (Qiagen, USA) for the isolation of genomic DNA from filtered water samples and algal mats samples following the manufacturers protocols. The PCR product from the extracted DNA samples were sent to NovoGene (NovoGene Ltd., Cambridge, UK) for Illumina sequencing on a MiSeq platform using Arch2A519F (CAGCMGCCGCGGTAA) and Arch1041R (GGCCATGCACCWCCTCTC) primers (Bižić et al., 2020a). Primers used from NovoGene (“16S/18S/ITS Amplicon Metagenomic Sequencing,” n.d.), included for Bacterial 16S V3V4 region are 341F (CCTAYGGGRBGCASCAG) and 806R (GGACTACNNGGGTATCTAAT), for archaeal 16S V4V5 region primers are Arch519F (CAGCCGCCGCGGTAA) and Arch915R (GTGCTCCCCCGCCAATTCCT), and for Eukaryote 18S V4 region primers are 528F (GCGGTAATTCCAGCTCCAA) and 706R (AATCCRAGAATTTCACCTCT). The PCR process was used to amplify the DNA to target the mcrA gene using the primer pair mlas-mod forward (GGYGGTGTMGGDTTCACMCARTA) (Steinberg and Regan, 2009) and mcrA-reverse (CGTTCATBGCGTAGTTVGGRTAGT) (Steinberg and Regan, 2009) using the Hot Star Taq Plus Master Mix Kit (Qiagen, USA) under the following conditions: 94°C for 3 min, followed by 30 cycles of 94°C for 30 s, 56°C for 40 s, and 72°C for 1 min, after which a final elongation step at 72°C for 5 min was performed. To check the success of the amplification and the relative intensity of the PCR product bands, a 2% agarose gel had been used for the mcrA gene. The libraries prepared using 25 ng of DNA samples.

To extract RNA from collected water samples and algal mat samples, RNeasy Power Water kit (Qiagen, USA) for the isolation genomic RNA. Before synthesis of first-strand cDNA, the DNA removal was done using the TURBO DNA-free Kit (catalog number AM1907, Thermo Fisher Scientific). The cDNA was synthesized using reverse transcriptase and incubated at 25 ⁰C for 10 min to extend the primers to the 42°C synthesis step, followed by placing the tube in a temperature-controlled thermal block at 42°C and incubate the reaction for 60 min. cDNA synthesis was terminated by incubating the reaction mixture at 70 ⁰C for 15 min. The cDNA was quantified using the Qubit RNA Assay kit (life Technologies), and RT-qPCR.

### Real time qPCR

To discern the levels of gene expression in our RNA samples, we opted for real-time amplifying qPCR due to its precision and speed. This approach employs fluorescent molecules to trace the progression of amplification as it occurs. The emitted fluorescence during this process is measured at each cycle, and this information is then employed to construct an amplification curve. Through a comparison between the fluorescence of the sample and established standards, we can reliably deduce the initial quantity of the target molecule within the sample. For 16S rRNA gene primers, we used 515 Forward (GTGYCAGCMGCCGCGGTAA) (Parada et al., 2016), and 805 Reverse (GACTACHVGGGGTATCTAATCC) (Herlemann et al., 2011). To detect methane producing organisms, we used mcrA gene, which encodes the alpha subunit of methyl-coenzyme M reductase (MCR), which catalyzes the final step in methanogenesis. It is present in all methanogens, making it a unique and universal functional marker for these organisms (Steinberg and Regan, 2009). We used Q5 High-Fidelity in master mix with Forward primer: mlas-mod-forward (GGYGGTGTMGGDTTCACMCARTA), Reverse primer: “mcrA-reverse (CGTTCATBGCGTAGTTVGGRTAGT) (Steinberg and Regan, 2009), templet DNA and water. The qPCR program included 35 cycles of denaturation in (95°C) for 30 s initial denaturation for 5 minutes, followed by annealing (60 °C) for 45 s, and ending with elongation (72 °C)for 45 s (Bižić et al., 2020b).

The success and relative intensity of the qPCR amplification products were analyzed using gel electrophoresis. This was performed on a 1.5% agarose gel containing SYBR Safe dye (diluted 1:10,000 in TAE buffer).

### DNA and cDNA sequencing and analysis

DNA samples were sent to NovoGene (NovoGene Co, Ltd., UK) for Illumina amplicon sequencing. The sequencing included 79 samples, 19 of which were from algal mats (covering habitat A, B, and D), 31 from 10µm filters (covering habitat A, B, C, and D), and 29 from 0.22µm filters (covering habitat A, B, C, and D). The primers used are documented in the section “DNA and RNA extraction and sequencing” using primers from NovoGene (“16S/18S/ITS Amplicon Metagenomic Sequencing,” n.d.). Sequencing results were grouped into five separate groups: 1) Bacteria: 16S V3V4 primers, 0.22µm and 10µm filter samples; 2) Bacteria: 16S V3V4 primers, algal mats; 3) Archaea: 16S V4V5 primers, 0.22µm and 10µm filter samples; 4) Archaea: 16S V4V5 primers, algal mats; 5) Eukaryotes: 18S V4 primers, algal mats and 10µm filter samples.

## Bioinformatics pipeline

The processing steps in the bioinformatic pipeline for this analysis was based on the DADA2 Big Data processing description (https://benjjneb.github.io/dada2/bigdata.html), as well as the tutorial steps for version 1.16 of the dada2 R package (https://benjjneb.github.io/dada2/tutorial.html). The implementation used the dada2 package installation from BioConductor 3.17 and R 4.3. Pipeline execution steps were defined as executable R scripts running in a Docker container image based on the official BioConductor docker image for BioConductor 3.17 with R 4.3. The underlying virtual machine running the docker container was an AWS EC2 t4g.2xlarge machine (8 VCPU, 32 GB RAM, 30 GB disk). Each pipeline execution was performed separately on each of the five groups above. The pipeline loaded data from NovoGene FASTQ files, which had already been merged and filtered (filenames ***sampleID***.extendedFrags.fastq.gz). Thus, the filtering and trimming steps of the pipeline did not do any additional trimming and truncating of the input data, which are the default settings. Only maxEE (max expected errors) were explicitly set to 1, which is lower than the default at 2 used by the dada2 software package. Steps to learn error rates, inter sequence variants and remove chimeras were done according to the Big Data pipeline description. The following assign taxonomy step used SILVA database nr 99 138.1 data set for 16S primer data, and PR2 v4.14.0 SSU DADA2 data for 18S primer data. The resulting sequence table data were saved for later additional analysis with R, using the R saveRDS() function. This allows data to be stored in a compact format that can easily be loaded into other analysis programs written in R.

### Amplicon sequence data analysis and visualization

The analysis and visualizations were created using the R phyloseq package, in combination with packages tidyverse, ggplot2, ggalt, ggsci, readxl, gt. Output data from DADA2 bioinformatics pipeline were combined with additional sample metadata, qPCR analysis data for RNA, flux and environmental forcing parameters from (Lundevall-Zara et al., 2021). These were combined into R phyloseq objects based on steps outlined in the DADA2 Pipeline Tutorial, section *Bonus: Handoff to Phyloseq*. (https://benjjneb.github.io/dada2/bigdata.html). Analysis and visualizations were done in RStudio bundled with the BioConductor 3.17/R 4.3 docker image, using Quarto Markdown (.qmd) notebook documents. Data from the bioinformatics pipeline were loaded as R RDS data using readRDS() function to produce phyloseq objects for each dataset from pipeline. For data processed with 16S V3V4 Bacteria primer, results for taxonomy Kingdom Archaea were excluded before further analysis, as these were not considered reliable with that primer. This was applied to the sample types algal mats, filter 0.22µm, and filter 10µm. For data processed with 16S V4V5 primer, only results for taxonomy Kingdom Archaea were included. Filtering out “methanogenic archaea” from this dataset was done by only including data whose taxonomy rank (Phylum, Class, Order, Family, Genus) included “Methan” in its name. This was applied to the sample types of algal mats, filter 0.22µm, and filter 10µm. Data processing also included calculating relative abundance for all reads in datasets, filter out low abundance ASVs, for 16S V3V4, keep ASVs > 0.5%, for 16S V4V5, keep ASVs > 0.1% and 18S V4, keep ASVs > 0.5%.

## Statistical analysis

PERMANOVA tests were run on datasets, using relative abundance at genus level, for the different primers and sample types to establish to what extent the variance could be explained by sampling period differences. Distances were calculated using Bray-Curtis. The R package vegan and its function adonis2() were used for the calculation.

NMDS analysis was done to see to what extent there may be taxonomic similarities between samples, based on habitat and sampling period. The NMDS ordination plots takes the calculated relative abundance data from the ASVs (with low abundance ASVS filtered out) and reduces the dissimilarities between ASVs from each of the samples into a 2-dimensional visualization, where points in the plot represent individual samples. The Bray-Curtis dissimilarity distance (Bray and Curtis, 1957) is calculated in the NMDS analysis to determine the similarity between the ASV datasets for each sample. A smaller distance between individual samples indicates that the ASV datasets more similarities in terms of relative abundance of the ASVs that are part of the samples, while larger distances indicate less similarity between samples – 0 means two datasets are identical, and 1 means no similarities at all. For each data set with relative abundance calculated and low abundance ASVs filtered out according to criteria in Data Tidying and Preparation section above. The ordination plot functionality of the R phyloseq package was used to produce the ordination plots.

## Results

### Spatial and temporal flux variations

Habitat A, characterized by dense vegetation, organic-rich sediment, and thick layers of floating algal mats on the water surface, exhibited the highest methane fluxes and concentrations throughout the study period (Figure 2). Methane fluxes in habitat A were lowest in June and October (approximately 1 mg m⁻² d⁻¹), with peak values of approximately 15 mg m⁻² d⁻¹ in July. Dissolved methane concentrations in habitat A varied from ∼50 μmol L⁻¹ in June to exceptional values exceeding 300 μmol L⁻¹ in July and September.

**Figure 2.**
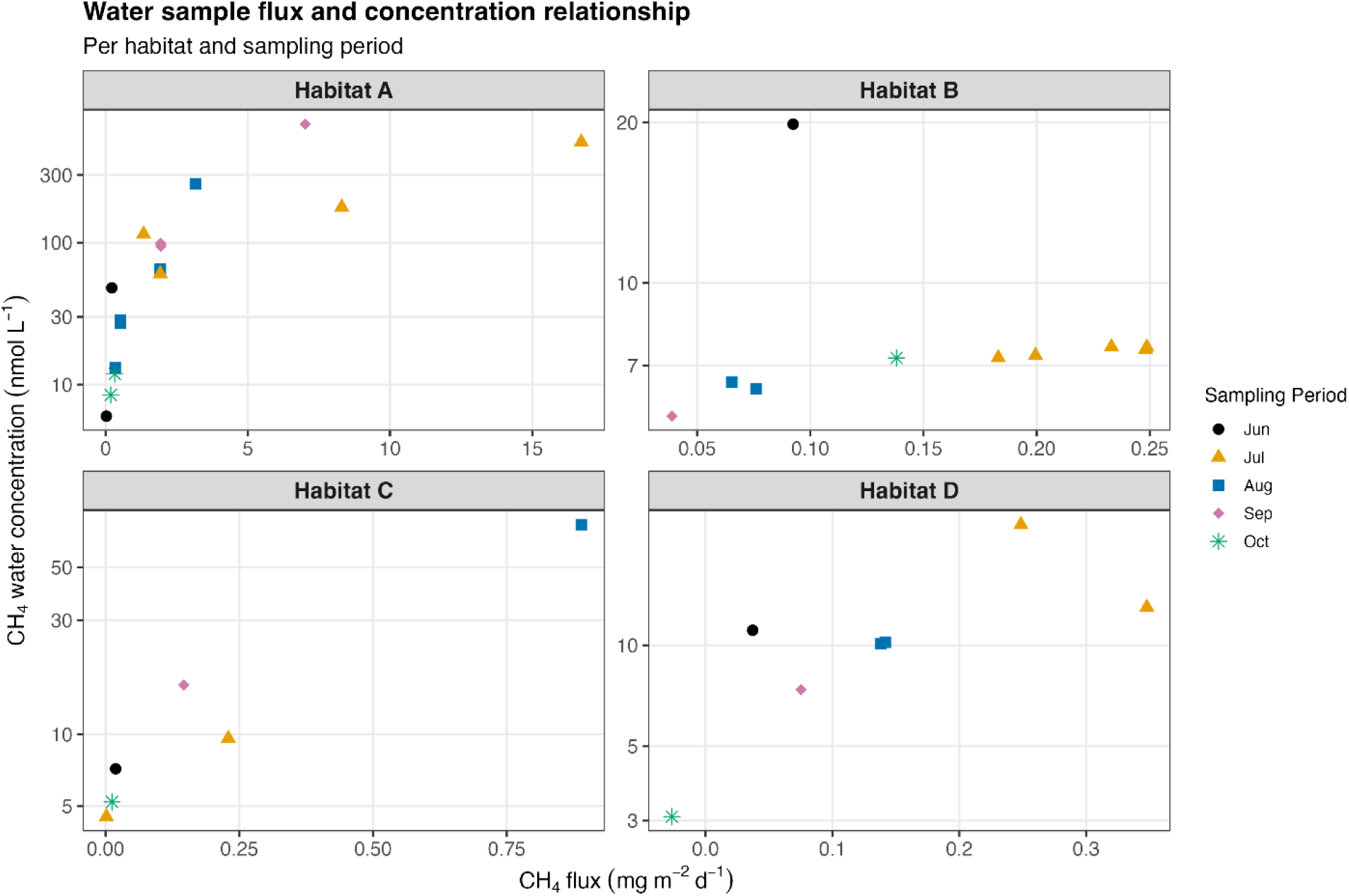
Flux and concentration of methane in water samples across all habitats and sampling periods, concentration on x axis and flux on y axis, with individual scales for each habitat. Sampling periods distinguished by color and shape of dot; each dot is a sample. Dashed trend lines connect multiple samples from the same sampling period. Concentration and flux for habitat A operate with a significantly larger variation than the other 3 habitats.

In contrast, habitats B, C, and D showed consistently lower and more stable methane emissions. Habitat B displayed moderate fluxes ranging from 0.05 to 0.25 mg m⁻² d⁻¹, with concentrations between 5-8 μmol L⁻¹ for most values across all sampling periods. Habitat C had fluxes ranging from 0.05 to 0.75 mg m⁻² d⁻¹ and concentrations from 5-70 μmol L⁻¹, peaking in August. Habitat D exhibited intermediate variability, with fluxes from 0.05 to 0.35 mg m⁻² d⁻¹ and concentrations ranging from 3-25 μmol L⁻¹.

The spatial gradient in emissions was pronounced, with habitat A fluxes being 20-60 times higher than other habitats during peak periods. Notably, all habitats except A showed relatively consistent flux-concentration relationships, while habitat A displayed extreme variability, particularly during the July sampling period when concentrations reached their maximum values. These spatial and temporal patterns remained consistent with results from the same coastal areas measured during 2019(Lundevall-Zara et al., 2021).

### Taxonomic classification based on phototrophic eukaryotes on 18S V4 at phylum level

Algal mat samples were collected from habitats A, B, and D. There were no algal mats in habitat C. The dominating classes were Metazoa, Dinoflagellata, Ciliophora, Ochrophyta, Fungi, and Chlorophyta in all habitats and sampling periods. Between 98.2 and 99.1 % of the 18S V4 sequences belonged to heterotrophs, with many representatives from Metazoa, Dinoflagellata, and Ciliophora being heterotrophic, and Fungi being entirely heterotrophic. These predominantly heterotrophic phyla were filtered away to display the relative abundances of known photoautotrophic eukaryotes only. In the algal mats Picochlorum constituted the major portion of the photoautotrophic community across sampling periods and showed some temporal fluctuation (Figure 3A). Additional taxa represented in Chlorophyta and Cryptophytae were consistently present, indicating a stable core photoautotrophic community despite some temporal changes.

**Figure 3.**
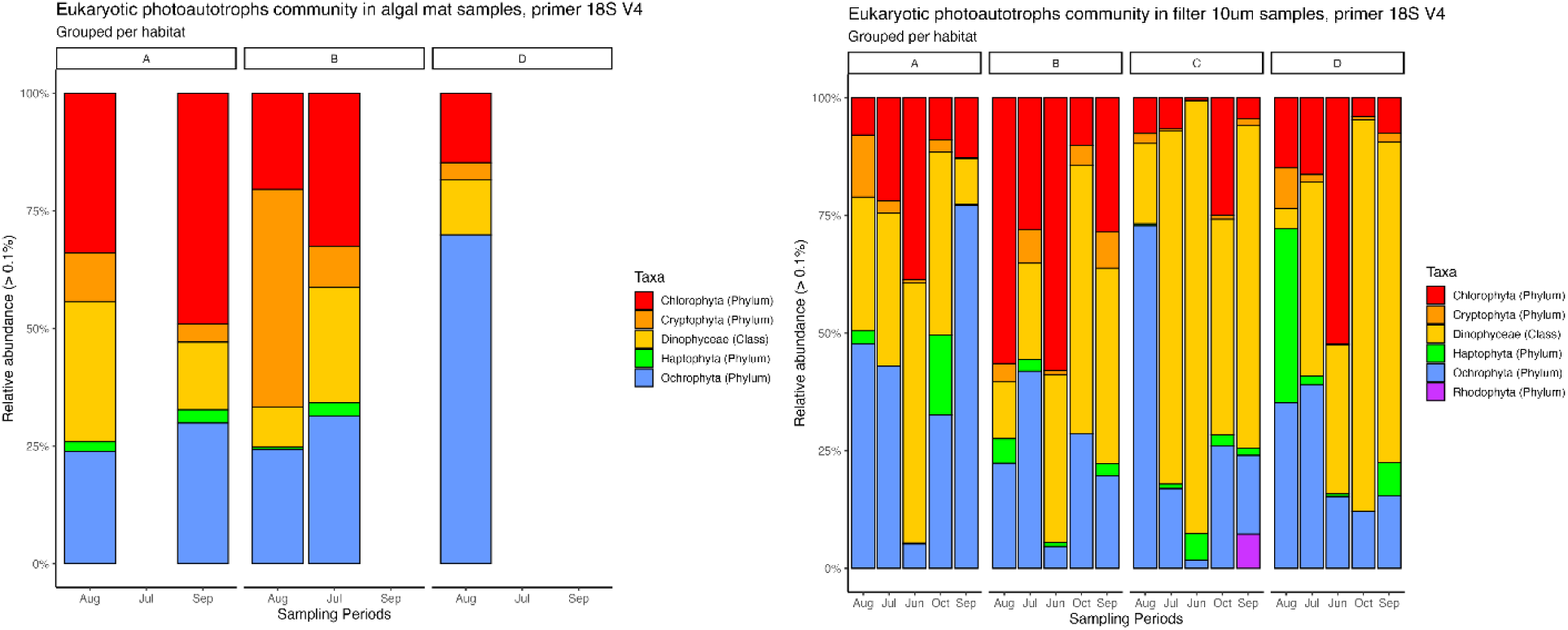
Average relative abundance for photoautotrophic eukaryote in eukaryotic community across habitat and sampling periods with 18S primer V4.

In the 10μm filtered samples, the distribution of taxa was more equitable, with multiple groups at similar abundance levels (Figure 5B). Temporal dynamics were more pronounced compared to algal mats exhibiting clear compositional shifts suggesting that planktonic communities were more responsive to seasonal variability compared to their algal mat counterparts (Dugenne et al., 2024; Miyagishima, 2023). Chlorophyta dominated in habitat A, B, and D during Juni, Juli and Augusti, while in habitat C the highest relative abundance occurred during October in filtered water samples. Metazoa and Ochrophyta were similar during all sampling periods across all habitats. Dinoflagellata and Ciliophora were higher in habitat C and D compared to habitat A and B during all sampling periods across all habitats.

A notable observation in Figure 3B is the apparent inverse relationships between certain taxa, where abundance increased in some groups correspond with decreases in others. This pattern might reflect competitive interactions or differential responses to environmental variables (Battistuzzi et al., 2023). Despite the shifting proportions of individual taxa, Figure 5B demonstrates consistent overall diversity levels across time.

The contrasting community structures between Figure 3A and Figure 3B point to distinct ecological niches and environmental influences shaping these photoautotrophic communities. The more stable, less diverse community shown in the algal mats likely reflects the structured nature of algal mat habitats, while the enhanced diversity displayed in the 10 um fraction suggests adaptation to a more heterogeneous planktonic environment (Iwai et al., 2024).

### Taxonomic classification of 16S V3V4 Bacteria

#### Algal mats

Taxonomic classification of bacteria was done to the phylum level. Only amplicon sequence variants whose relative abundance is >0.5% of prokaryotic community were considered for the analysis. In algal mats the bacterial community was dominated by Proteobacteria, Bacteroidota, Cyanobacteria, Verrucomicrobiota, Firmicutes, and Patescibacteria (Figure 4a). The relative abundance of Desulfobacterota in our samples was less than 0.7% in algal mat samples (Figure 4-a).

**Figure 4.**
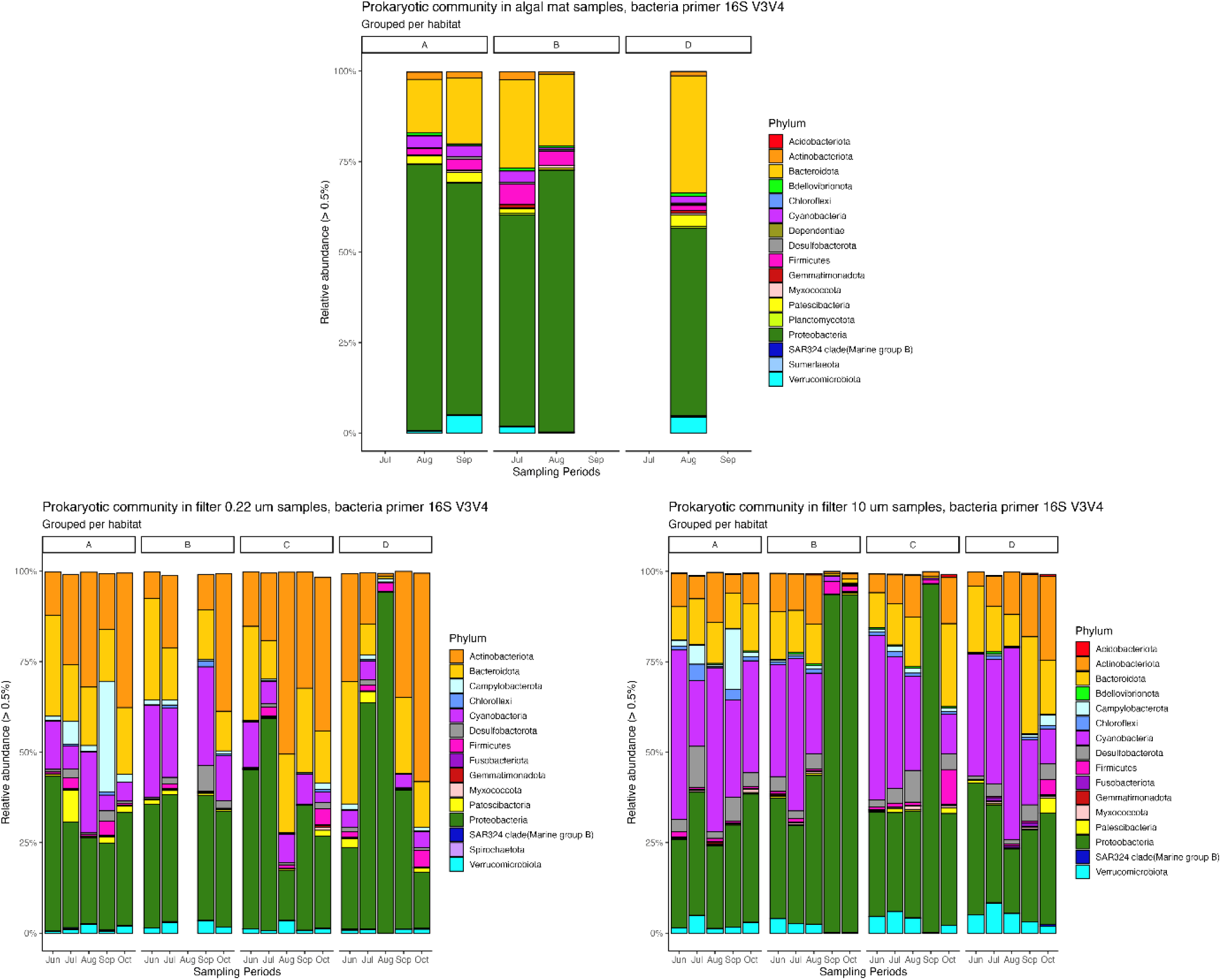
Prokaryotic community at phylum level grouped by habitat and sampling period with bacteria primers 16S V3V4 for: a-Algal mat samples, b-Filter samples 10 µm, c-Filter samples 0.22 µm.

**Figure 5.**
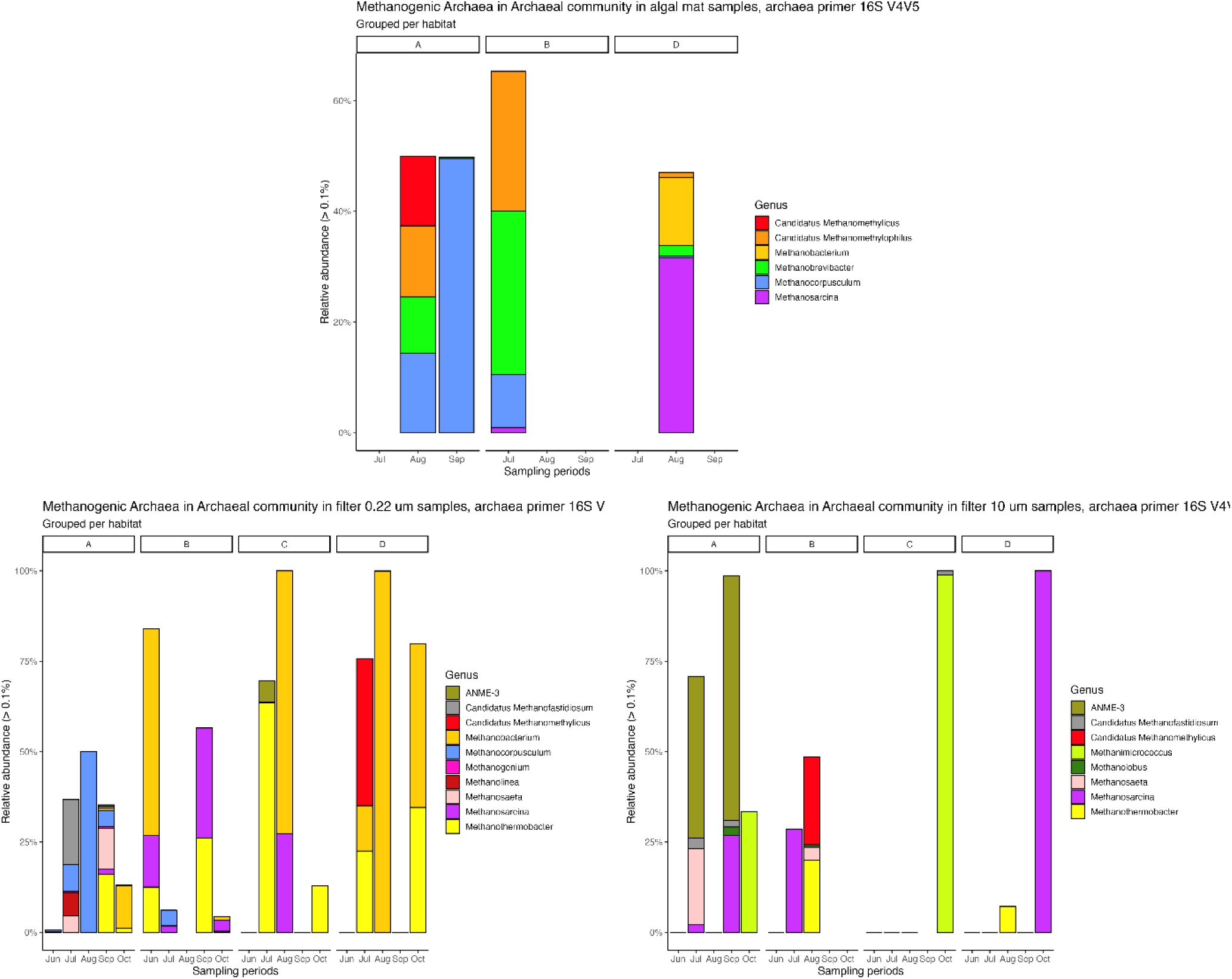
Average relative abundance for methanogenic archaea in archaeal community across habitat and sampling periods with archaea primers V4V5. The taxonomic classification is displayed at the genus level for: a-Algal mat samples, b-Filter samples 10 µm, c-Filter samples 0.22 µm.

#### 0.22 μm fraction

Filtered water samples of the 0.22µm size fraction were dominated by Actinobacteriota, Bacteroidota, Cyanobacteria, Proteobacteria, and Verrucomicrobiota. Proteobacteria dominated from June to October, showing particularly high and stable proportions in habitat A (25-45% relative abundance), while being more variable in habitats B, C, and D where they ranged from 0-90% depending on sampling period. Actinobacteria and Bacteroidetes displayed a distinct seasonal variation with peak abundances during the late summer months of August and September. Planctomycete were most prominent in the 0.22 μm fraction, although their abundance was relatively low showing less seasonal variation compared to the other phyla. Anaerobic, sulfate-reducing bacteria belonging to the phylum Desulfobacterota, specifically Desulfobulbus clades, were detected in the amplicon sequences of both 10 µm and 0.22 µm filtered water samples. Notably, in the 0.22 µm filtered water samples, we also identified bacteria of the phylum Desulfobacterota. The relative abundance of Desulfobacterota in our samples was less than 7.0% in 0.22 µm filtered samples (Figure 4-b, c). Vibrio strains made up less than 0.5% of all bacterial sequences in all sample types and habitats.

#### 10 μm fraction

Actinobacteriota, Bacteroidota, Cyanobacteria, Proteobacteria, and Verrucomicrobiota. Actinobacteria and Bacteroidetes were most abundant in this fraction and most prominent in habitats B and C during August. In all sample types, we identified cyanobacteria sequences, but most were found in the >10 µm size fraction. Cyanobacteria showed clear temporal patterns with higher abundance in June, July and August, and their peak distribution varied considerably between habitats, with habitat A between approximately between 20-50% being the most stable temporal distribution, and habitat B the most varying with approximately 0-50%. Proteobacteria observed the highest peaks in habitat B, and C with peaks at approximately 90% relative abundance. The relative abundance of Desulfobacterota in our samples was less than 11.2% in 10 µm filtered samples (Figure 4-b).

### Taxonomic classification in 16S V4V5 Archaea at genus level

Figure 5 illustrates the distribution of methanogenic archaea within the archaeal community for the different habitats and sampling periods. The dominant archaea at the genus level varies for different sampling periods within each habitat, and for different sample types. All the methanogenic archaea belong to the phylum Euryarchaeota. For algal mats in habitat A (as depicted in Figure 5-a) Candidatus Methanomethylicus, Candidatus Methanomethylophilus, Methanobrevibacter, and Methanocorpusculum dominated during the August sampling period. Notably, Candidatus Methanocorpusculum dominated methanogenic archaea in habitat A during august, when it represented close to 100% of all detected methanogenic archaea (50% of archaeal community), but no more than 9.6% and 0.3% of the archaeal community in samples from habitats B and D, respectively.

In habitat B, the dominant genera were Candidatus Methanomethylophilus, Methanobrevibacter, Methanocorpusculum, and Methanosarcina. Conversely, in habitat D, the average relative abundance at the genus level pointed towards the dominance of Methanosarcina and Methanobacterium (order Methanobacteriales) for algal mats (Figure 5-a). For Methanosarciniales we detected the genus Methanosarcina in habitats B (July) and D (August), with relative abundance up to 67.2% of methanogenic archaea (31.6% of archaeal community) in habitat D, and up to 1.3% of methanogenic archaea (0.9% of archaeal community) in habitat B (Figure 5-a).

Methanosarciniales and Methanomassiliicoccales are present only in the algal mat samples in our data, while Methanobacteriales is present in all sample types (Figure 5-a, b, c). Methanobacteriales, including Methanobacterium, were detected in all habitats in filter 0.22 µm samples (Figure 5-c). Also, Methanomicrobiales were detected in filter samples with a pore size of 0.22 µm from the same habitats and sampling periods (Figure 5-c). We also detected Methanosarcina in filter samples (Figure 5-b, c).

A high abundance of sequences indicating anaerobic methanotrophy by ANME-3 was found in the 10-micron fraction and correlates with the presence of Desulfobulbus sequences, supporting the presence of consortia that can carry out anaerobic oxidation of methane by sulfate or sulfur reduction (Knittel and Ravenschlag, 2019; (Knittel and Boetius, 2009; Milucka et al., 2012)Milucka et al.)

While DMS-based methylotrophy occurs in both sediments and the water column due to DMSP degradation from diatoms, the co-occurrence of Methanolobus with sediment-associated taxa such as ANME-3 and Desulfobulbus suggests that the observed signal likely originates from sediment-driven anaerobic DMS metabolism, as supported by recent findings for Baltic Sea sediments (Tsola et al., 2024).

Methanolobus is present in habitat A (≤6%) only in filtered water samples 10 µm in September, and in habitat B (≤0.7%), only in filtered water samples 10 µm, and only in August. Methanolobus, ANME-3 and Desulfobulbus only appeared together in habitat A during September.

Analysis of the methanogenic pathways based on the relative abundance patterns reveals interesting temporal dynamics across the different sampling conditions. This pattern shifts in September, where we observe increased community diversity and a notable rise in Methanosarcina abundance.

In the filter 0.22 µm samples, the community structure appears more dynamic across sampling periods. The substantial presence of Methanosarcina suggests multiple pathways are active simultaneously.

The filter 10 µm samples display less diverse communities but exhibit pronounced temporal shifts. Early sampling periods show a mixed community with significant Methanosarcina presence, indicating multiple active pathways (Chen et al., 2017; Mori et al., 2012). Later periods demonstrate strong dominance by either Methanobacterium or Methanosarcina.

Throughout all samples, hydrogenotrophic methanogenesis appears to maintain a consistent presence, indicated by the persistent detection of Methanobacterium.

At genus level across different habitats and sampling periods in 10 µm filter samples methanogenic Archaea. Are dominated by *Methanosarcina* dominated at habitat A, B and D during the warm sampling periods at July, August, and September, while Methanimicrococcus dominated during the fall, October-sampling-period at habitat B and C (Figure 5B).

In 10 µm filtered water samples, our study found the presence of the Pseudomonas genus in all sample types, with a significant presence in August. Pseudomonas is most abundant in samples with a 0.22 µm filter. Previous research by Wang et al., (2017) demonstrated that the Pseudomonas group can efficiently use methylphosphonates for methane production.

Figure 5-c shows the Methanogenic Archaea community at genus level grouped by habitat and sampling period in 0.22µm filter samples for Archaea primers 16S V4V5 amplicon sequence. The Methanogenic Archaea dominated in habitat A, are Methanothermobacter, Methanocorpusculum, Methanolinea, Methanosaeta, Methanosarcina, Methanobacterium, and Candidatus Methanofastidiosum (Figure 5-c). While in habitat B dominated Methanothermobacter, Methanocorpusculum, Methanosarcina, and Methanobacterium (Figure 5-c). The Methanogenic Archaea dominated in habitat C, are Methanothermobacter, ANME-3, Methanosarcina, and Methanobacterium. In habitat D the dominated Archaea in genus level reads Methanothermobacter, Methanobacterium, Methanosarcina, and Candidatus Methanomethylicus.

Looking at the temporal variation across the samples, there are distinct shifts in the methanogenic archaeal community composition throughout the different months. This temporal dynamism suggests strong seasonal influences on these microbial populations (Fu et al., 2023). For instance, Methanobacterium shows particularly strong representation during the July-August period.

The presence and abundance of dominant genera provide interesting insights into community structure. Methanobacterium, Methanocella, and Methanosarcina emerge as key players, but their relative abundances fluctuate considerably across samples.

Habitat specificity is another notable pattern, with different sampling locations showing distinct methanogenic communities. The algal mat samples, in particular, show unique community profiles compared to the filtered samples.

### Taxonomic community correlation analysis

Picoplankton, defined as organisms smaller than 3 micrometers, encompasses both freshwater picocyanobacteria (including *Synechococcus, Synechocystis,* and *Cyanobium*) and marine picocyanobacteria (predominantly *Synechococcus* and *Prochlorococcus*). These microorganisms play a pivotal role in aquatic methane production, as demonstrated by recent studies (Fazi et al., 2021; Tenorio and Farías, 2024). The findings indicate that picoplankton are significant contributors to methane production, particularly in the presence of methylated substrates such as methylphosphonic acid and trimethylamine (Tenorio and Farías, 2024). Methane production involving picoplankton, including heterotrophic bacteria and cyanobacteria, is integral to the recycling of dissolved organic carbon and the facilitation of methylotrophic methanogenesis (Tenorio and Farías, 2024).

In our study, all four aforementioned Picocyanobacteria were identified, though only *Cyanobium* exhibited a relative abundance exceeding 1% in any combination of habitat and sampling period. Notably, relative abundances above 10% were predominantly observed in habitat B, with a peak value of 17% recorded in July. In contrast, the relative abundances of the other three Picocyanobacteria (*Synechococcus, Synechocystis,* and *Prochlorococcus*) were generally below 0.2% across all habitats and sampling periods. Picocyanobacteria were detected in all habitats and sampling periods in filtered water samples, while in algal mat samples, they were primarily found in habitats A, B, and D, particularly during July and August.

### NMDS analysis

The purpose of the NMDS analysis is to get an understanding if there are any discernable similarities in terms of microorganism composition for the different samples, in relation to the sample types, as well as the spatial and temporal parameters for the samples. The NMDS analysis for bacterial primers included all bacteria in Figure 6. For archaea, the focus of the NMDS analysis was on methanogenic archaea (Figure 7), and for eukaryote, the focus was on photoautotrophic eukaryote (Figure 8). The NMDS plots point to notable differences between habitats, with habitat A seeing more distinct clustering effects for bacteria and methanogenic archaea (Figure 6 and Figure 7). However, the number of samples are overall fewer for habitats B, C, and D, for bacteria and methanogenic archaea, which may contribute to less distinguishable groups based on sample type. Clustering samples based on sampling period can limited extent be observed in habitat A.

**Figure 6.**
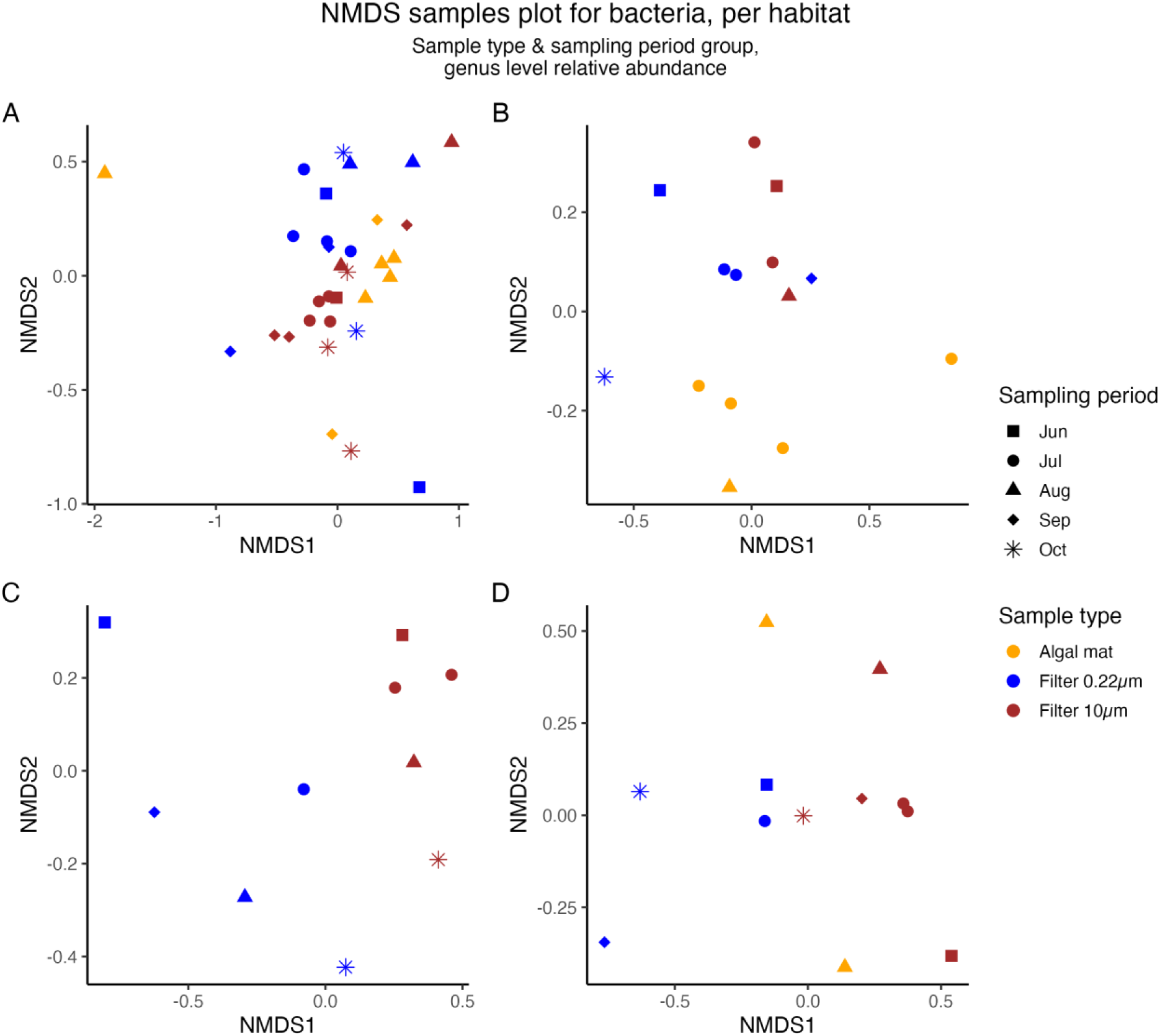
NMDS analysis for Bacteria, with separate diagrams per habitat and plotting sample relationships. Samples are color coded based on sample type (algal mat samples, filter 0.22µm samples and filter 10µm samples), and separate shapes for different sampling periods. For each sample type there are for the most part notable clustering patterns, and more separated samples in habitats B, C, and D, while habitat A shows closer ordering between samples, also across sample types.

**Figure 7.**
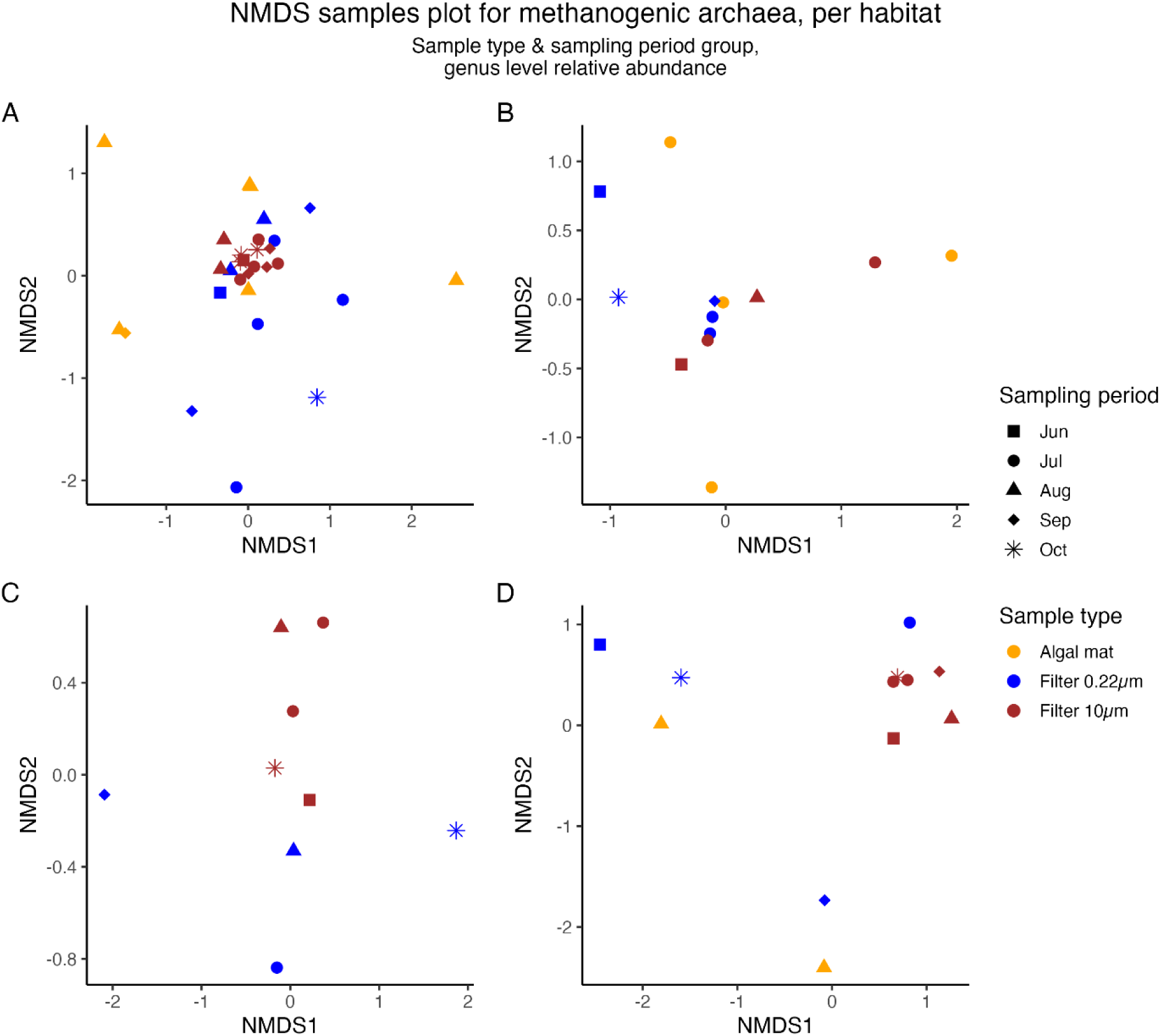
NMDS analysis for methanogenic Archaea with separate diagrams per habitat and plotting sample relationships. Samples are color coded based on sample type (algal mat samples, filter 0.22µm samples and filter 10µm samples), and separate shapes for different sampling periods. There are distinct differences in the ordering of 0.22 and 10 µm fractions, suggesting that variations of methanogenic archaea involve other contributions than potential sediment resuspension.

**Figure 8.**
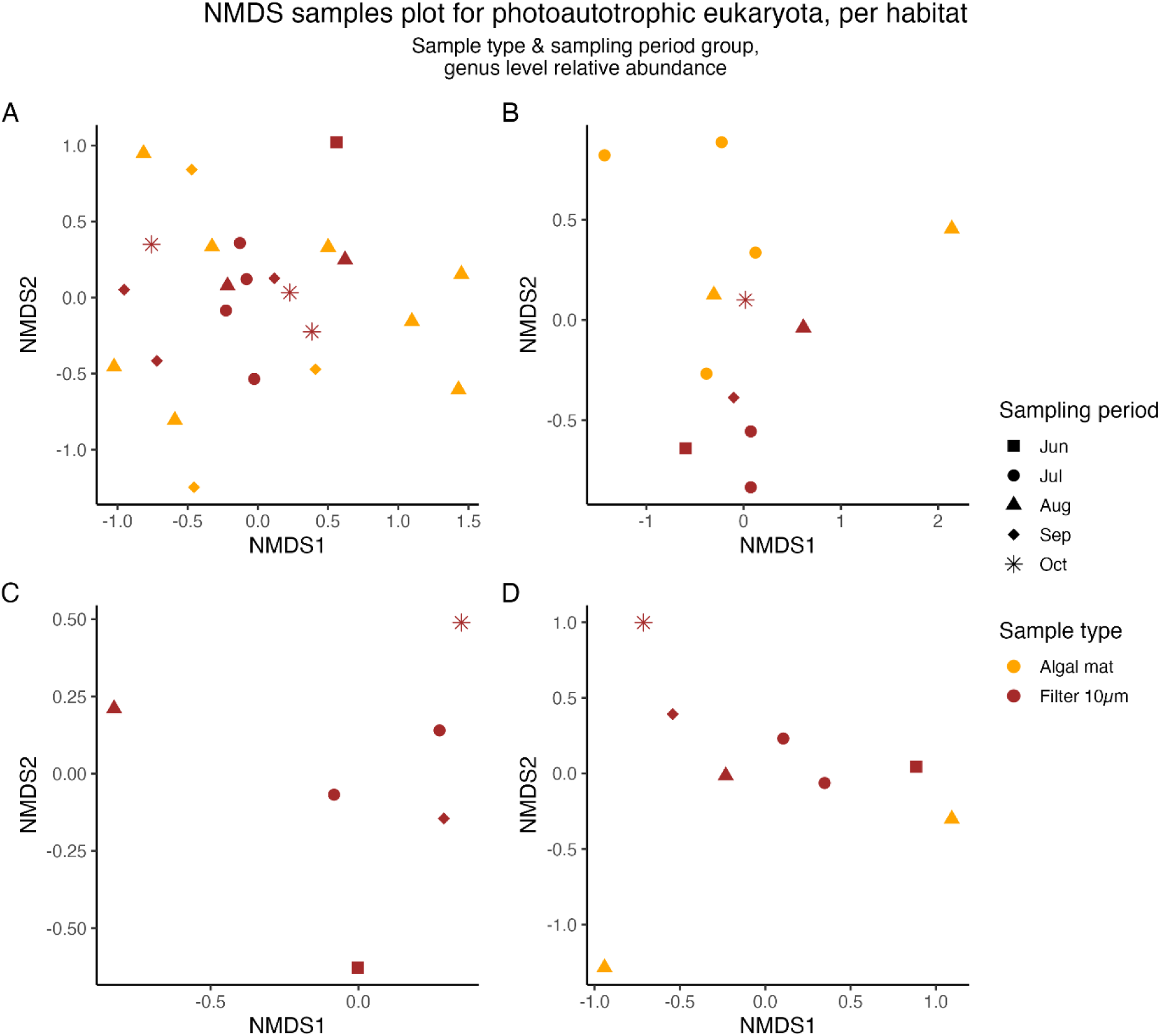
NMDS analysis for photoautotrophic Eukaryote with separate diagrams per habitat and plotting sample relationships. Samples are color coded based on sample type (algal mat samples, and filter 10µm samples), and separate shapes for different sampling periods. The distribution of samples supports what is shown in Figure 3, with similar taxonomic composition although different proportions.

### Real time amplifying qPCR in RNA samples

The Delta CT levels are an indication of gene expression levels of mRNA molecules. To establish the levels of gene expression in our RNA samples, real-time qPCR was used. Expression levels are shown as Delta CT (Mean RT-qPCR mcrA CT-Mean RT-qPCR 16S CT). All habitats with algal mat samples showed positive mcrA-16S rRNA (Delta Ct), but the Delta Ct is significantly lower for algal mat than for the filter samples (Figure 9). The filter samples all showed positive mcrA gene expression levels in all habitats for the pore size 10 µm filter, while there are only gene expressions indicated in habitats A, B, and D for pore size 0.22 µm.

**Figure 9:**
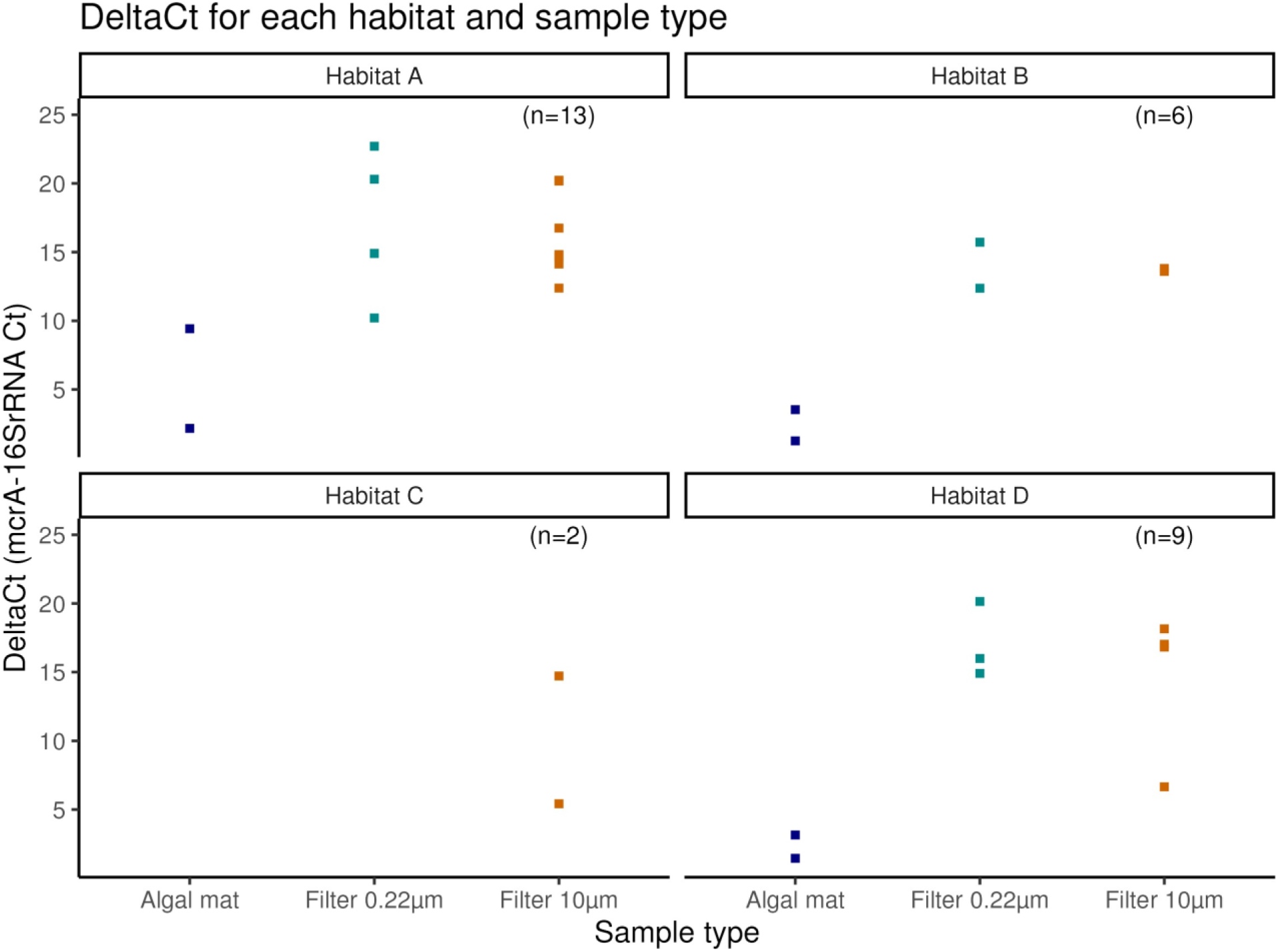
Results of qPCR analysis, delta Ct for each habitat and sampling type.

The results from real time amplification qPCR in our RNA samples (Figure 9), confirm that the average mcrA gene expression is significantly higher in habitat A compared to other habitats in all sample types. Delta Ct levels are similar in filtered water samples 0.22µm and 10µm samples, and lower for algal mat samples.

Habitat D, which has hard unvegetated ground, and is surrounded by rocky cliffs, has the second highest level of gene expression despite sparse sessile aquatic vegetation and algal mats during summer. Gene expression levels were relatively lower in habitat B and C.

### Statistical analysis

For both V4V5 Archaea primer and V3V4 Bacteria datasets, the analysis indicated that there were significant differences between the 10 and 0.22 μm filter datasets for both the archaeal and bacterial communities, while the algal mat dataset did not show significant differences between sampling periods (Table 2).The number of sampling periods included for the algal mat dataset were fewer, and fewer data points than for the filter sample types. Thus, the results for algal mats are inconclusive in that regard. However, the PERMANOVA results agree with the NMDS analysis, which indicate differences between sampling periods for filter samples to a higher degree than for algal mats.

For the 10 and 0.22 μm filter samples the PERMANOVA tests show higher R-squared values than the test for the algal mats, indicating that sampling period explains more of the variation in Archaeal and Bacterial composition in the filter samples (Table 2). The R-squared values are also notably higher for the archaea filter datasets than for the bacteria filter datasets.

### Our study also investigated the presence and occurrence of iron, nitrate and sulfate reducers

The co-occurrence of such reducers may indicate varying redox conditions, which can occur in sediments (Jin et al., 2017). These groups also thrive in oxygen-limited and anaerobic environments (Aeppli et al., 2023; Sørensen, 1982), and often rely on organic matter as an electron donor, which may indicate organic carbon (Jin et al., 2017).

We picked seven reducers to analyze; Geobacter and Shewanella as iron reducers(Li et al., 2023), Acidovorax and Dechloromonas as nitrate reducers(Carlson et al., 2013; Liu et al., 2019), Desulfobacter, Desulfobulbus, and Desulfovibrio as sulfate reducers (Li et al., 2023). We also checked for manganese reducer ANME-2d, also known as Methanoperedenaceae (Leu et al., 2020), but these could not be found. The 16S community data contained overall low levels of these reducers; 98% of reducer values < 1% relative abundance, 83% of values < 0.30%, and 63% < 0.10%. Only Shewanella (max value 1.45%) and Dechloromonas (max value 0.70%) had some slightly higher relative abundance values compared to the others.

All three types of reducers (iron, nitrate, and sulfate) co-occurred only in habitat A in July and September in filtered water samples in relative abundances >0.10%, all other combinations of habitats and sampling periods had either zero, one or two types of these reducers. Algal mat samples had presence of all reducers, but notably higher levels of nitrate reducers in habitats A, B and D, almost all of these during August (0.14 – 0.31% relative abundance). Iron and sulfate reducers were < 0.04% relative abundance, if present.

In our study we try to shed light on the distribution and abundance of iron, nitrate, and sulfate reducers across a gradient of habitats characterized by varying vegetation density, sediment organic content, and water depth. The presence of all three types of reducers could indicate the presence of resuspended sediment, although with abundance levels far below 1%, this may also be just background noise. The overall low abundance of these reducers suggests that while they are present, they may play a minor role in the microbial community structure of the studied ecosystem. Habitat A, being the most vegetated and organic-rich area with shallow water depths (0.5 meters), supports a diverse community of reducers, as evidenced by the co-occurrence of all three types in filtered water samples during July and September. This finding highlights the importance of habitat characteristics, including vegetation and organic content, in fostering a conducive environment for microbial diversity, and could potentially have elements of sediment. In contrast, Habitat C, with deeper water (up to 2 meters), may present different ecological conditions that limit the abundance and diversity of reducers, as well as habitat D, with hard unvegetated ground. The absence of all three reducer types in these habitats during certain sampling periods suggests that depth and associated factors—such as light penetration, nutrient availability, and sediment characteristics—may significantly influence microbial community composition. The higher abundance of nitrate reducer bacteria in algal mats of shallow habitats A, B, and D, particularly during August, indicates a potential seasonal interaction between primary producers and microbial reducers. This relationship underscores the complex interplay between environmental forcing factors, habitat structure, and microbial activity, particularly in shallow, nutrient-rich areas. Overall, the presence of these reducers across all habitats, despite their low abundance, points to their potential roles in biogeochemical processes, such as nutrient cycling and energy flow, within these aquatic ecosystems, as well as potential methane production in oxic water surface. Future research should focus on analyzing the functional significance of these microbial groups, examining how water depth, vegetation, and seasonal variations influence their distribution and activity, and exploring the interactions between reducers and other microbial communities to gain a more comprehensive understanding of the ecosystem’s microbial ecology.

**Cyanobacteria** have been observed to produce methane as a secondary by-product, though the underlying mechanisms of this process remain incompletely understood. The levels of methane produced by cyanobacteria are significantly lower than those observed in methanogenic archaea. Methane production by cyanobacteria occurs under specific conditions, such as during the degradation of methylated compounds or as a byproduct of their metabolic processes (Zou et al., 2024). Methane-producing cyanobacterial strains, including *Nodularia* PCC 9350, *Synechocystis sp.* PCC 6803, and *Trichodesmium*, were detected in all sample types and sampling periods across various habitats. A comparative analysis between the occurrence of these cyanobacterial groups and methanogenic archaea revealed that these methane-producing cyanobacteria were consistently present across all sample types, sampling periods, and habitats (Figure 10). The strain *Nodularia* PCC 9350 has noticeably higher relative abundance compared to the other strains of cyanobacteria in the comparison. The fact that *Nodularia* PCC 9350 mostly detected in 10 µm filtered water samples means that it is present in free water column. This means ecologically that aquatic methane emission must account for methane production in surface water as well as methane emission caused by archaea in sediment.

**Figure 10.**
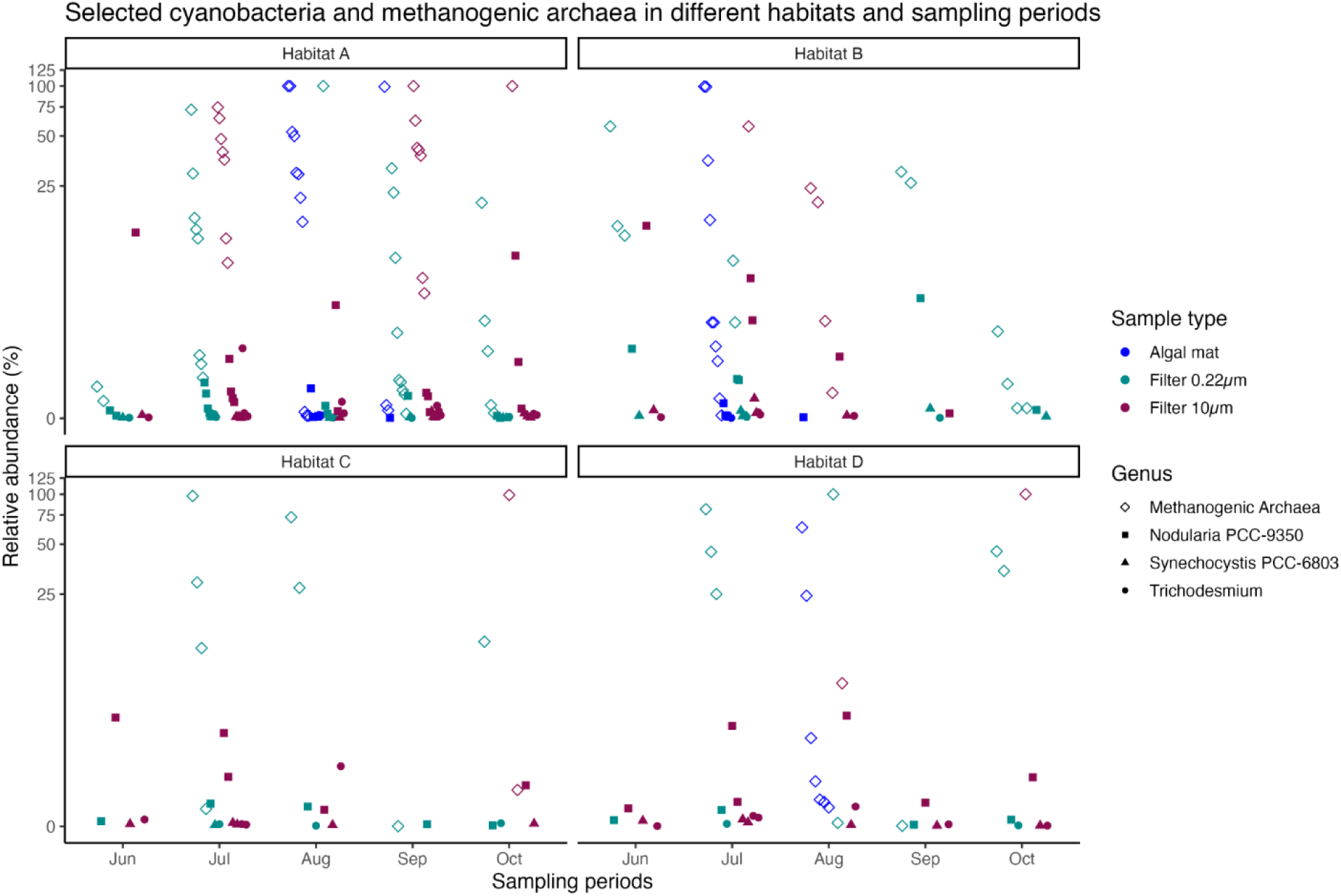
Selected cyanobacteria and methanogenic archaea across all habitats and sampling periods.

**Figure 11.**
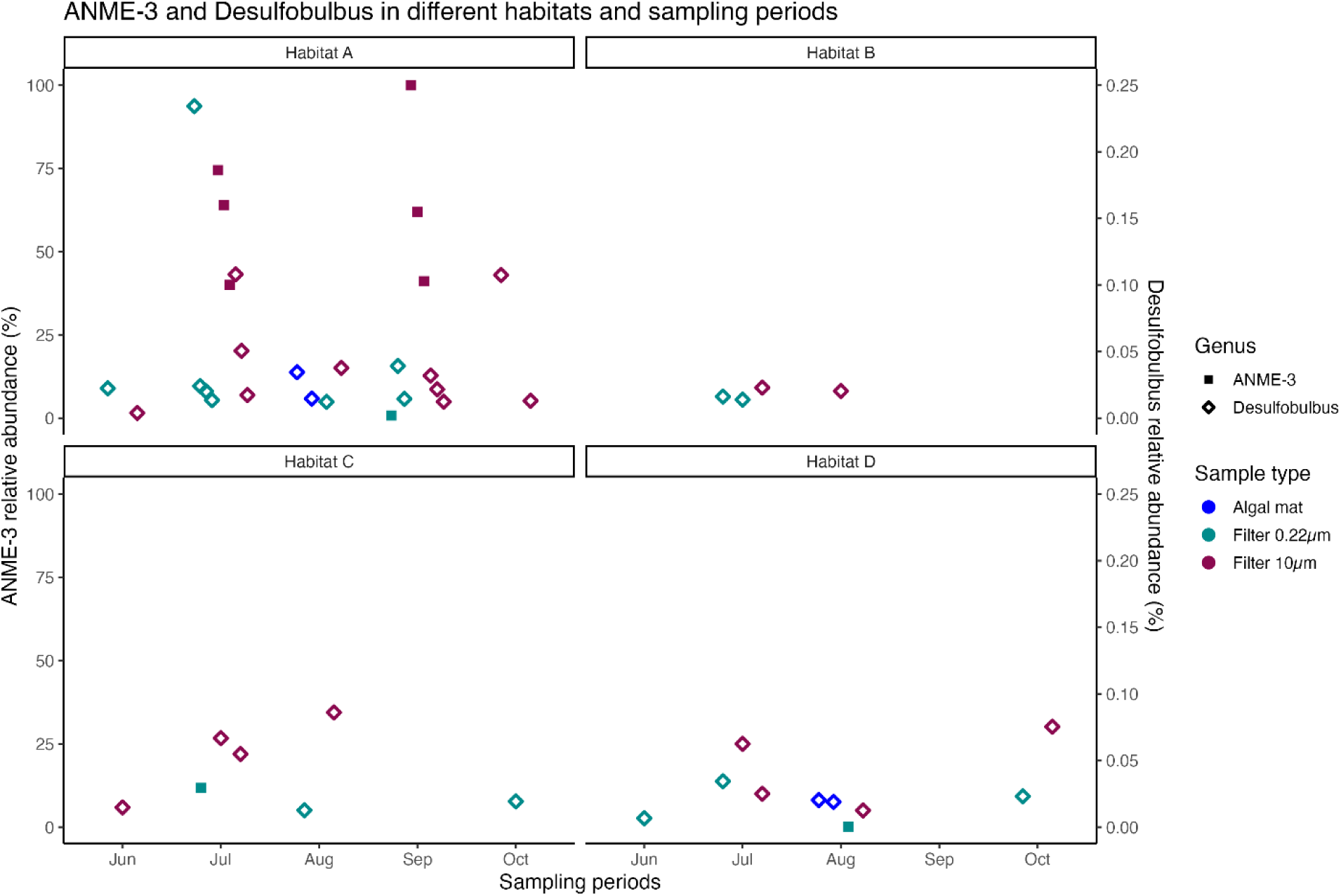
ANME-3 and Desulfobulbus in different habitats and sampling periods.

In shallow areas of the Baltic Sea, Nodularia (a cyanobacterium) and methanogens (anaerobic archaea) can interact but aren’t dependent on each other (Laamanen et al., 2001; Medwed et al., 2024). Their potential interactions are enhanced in shallow waters because decaying Nodularia blooms quickly create anoxic conditions near sediments and shallow depth allows more direct nutrient cycling between organisms (Medwed et al., 2024). Also, seasonal changes in these shallow waters can intensify their interactions (Medwed et al., 2024).

While these organisms can benefit from each other’s presence, they can and do exist independently in the Baltic Sea environment (Teikari et al., 2018b).

In algal mats, Nodularia and methanogens develop a closer relationship through the mat’s layered structure, which may create conditions for both organisms – aerobic zones for Nodularia’s photosynthesis at the top and anaerobic zones for methanogens below. This physical proximity enables direct metabolic exchange while providing stability and protection for both groups.

The presence of Synechocystic PCC-6803, which is a freshwater cyanobacterium, may indicate some adaptation to more brackish conditions, given that it is present in all habitats and sampling periods (Iijima et al., 2015). While the presence of Trichodesmium may indicate active nitrogen fixation (Karl et al., 2002). It is present in all habitats. The relative abundance levels for both Trichdesmium and Synechocystic PCC-6803 are low, and only Trichodesmium can be found at relative abundance levels above 0.5%, in habitat A and C and then in July and August, in habitats A, C and D.

In summary, our samples from both filtered water and algal mats are likely to show different patterns of association – the filtered water samples may show looser, more independent relationships, while the mat samples are likely to demonstrate stronger functional linkages and more structured interactions between Nodularia and methanogens due to their close physical proximity and shared microenvironment.

The comparison between relative abundance of ANME-3 and Desulfobulbas shows that the Desulfobulbas is represented in all sample types across all habitat and sampling periods, while ANME-3 is mostly represented in 10 μm filtered water samples in habitat A. ANME-3 also occurs in some of 0.22 μm filtered water samples in habitat A, C, and D, although at lower relative abundance than the 10 μm filtered water samples in habitat A.

The presence of high levels of ANME-3 in habitat A in July and September may potentially have been affected by sampling boat that may potentially have disturbed sediment, given the very shallow areas with vegetated ground in habitat A. At the depths in habitat C, the sampling boat is unlikely to have caused disturbance, and in habitat D the ground is unvegetated, also unlikely to have an effect from sampling boat. Thus, the presence of ANME-3 in 0.22 µm filtered water samples points to sources from the free water column.

Desulfobulbus is mainly represented in the filtered water samples, both in 10 µm and 0.22 µm, for most sampling periods, and in August in algal mats as well for habitats A and D. The latter might suggest oxygen-depleted zones within the algal mats.

The presence in the filtered water samples might suggest localized anaerobic or low-oxygen microenvironments, and the detection of Desulfobulbus in 0.22 µm filtered water samples suggest free-living planktonic bacteria in the water column (Diao et al., 2018; Giloteaux et al., 2013).

## Discussion

### Evidence for potential in-situ methane production in oxic Baltic Sea waters

RT-qPCR detection of mcrA and abundant reads of methanogenic Archaea in algal mats, on 10 µm filters and on 0.22 µm filters provide evidence for potential in-situ production of methane in the oxic nearshore waters of the Baltic Sea. These data add to the growing number of studies that indicate methane formation in oxic waters. Since the locus of formation is close to the sediment-water interface, these surface waters may comprise a direct emission source to the atmosphere.

In addition, studies have highlighted the role of microbial breakdown of methylphosphonate (MPn) in methane production within aquatic ecosystems. For example, MPn decomposition has been shown to contribute to methane emissions in surface oceans and lakes (Taenzer et al., 2020). This process is often linked to phosphate-limited conditions, which commonly coincide with methane-supersaturated waters(Mesquita et al., 2023; Tenorio and Farías, 2024).

Cyanobacteria have also been implicated in MPn-driven methane production, particularly under oligotrophic conditions (Bižić et al., 2020b). These findings suggest that multiple microbial pathways may contribute to methane cycling in aquatic environments, particularly in nutrient-depleted settings. High methane concentrations in oxic surface waters close to the seafloor boundary have been connected to the rapid upward movement of methane from bottom sediments. This phenomenon is particularly common in areas where the boundary between sulfate and methane zones lies close to the surface, often due to eutrophication or sediment buildup. Such circumstances can result in direct emissions of methane from surface waters into the atmosphere (Żygadłowska et al., 2024).

#### Potential sediment source of Archaea

A critical question emerges regarding whether the detection of Archaea and positive qPCR results for mcrA genes could be attributed to sediment suspension or bubble-mediated transport rather than in-situ production. Several lines of evidence address this possibility. The microbial community composition across the three size fractions (algal mats, 10 μm filters, and 0.22 μm filters) exhibits distinct characteristics and demonstrates temporal variability throughout the sampling period. The abundance of aerobic eukaryotic heterotrophs in the algal mats supports extensive oxygen consumption within these microenvironments, potentially creating anaerobic niches conducive to methanogenic activity.

Furthermore, the presence of sequences indicative of bacterial anaerobic heterotrophy, including sulfate reduction, iron reduction, manganese reduction, and anaerobic oxidation of methane, suggests the establishment of redox gradients within the mat structures.

Comparative analysis of these sequences with published sediment data from Broman et al., (2022) reveals both similarities and differences in microbial community composition, providing insights into the potential sedimentary versus in-situ origins of the observed methanogenic activity.

The distinct seasonal patterns in microbial community structure, combined with the metabolic diversity observed across different size fractions, suggest that while sediment resuspension may contribute to the observed archaeal presence, the evidence supports genuine in-situ methanogenic processes occurring within the oxic water column, particularly in association with algal mat microenvironments.

## What do the data tell us about the role of photosynthetic organisms in methane production?

The 10μm fraction captured larger colonial cyanobacteria and aggregates. These 10 µm fraction colonies engage in photosynthesis and generate particulate organic matter while creating oxygen gradients around their structures (Kihara et al., 2014; Mehdizadeh Allaf and Peerhossaini, 2022). The colonial formations potentially harbor localized anoxic microsites where attached methanogenic communities might thrive (Arvidson and Baker, n.d.). This fraction shows pronounced seasonal dynamics, with higher cyanobacterial abundance during warmer periods.

Floating algal mats, however, can create anoxic microenvironments conducive to methanogenesis. This is supported by micrometer-scale measurements of oxygen through algal mats and floating aggregates have provided direct evidence for the existence of anoxic conditions inside such layers despite oxygen-saturated conditions in the ambient water (de Angelis and Lee, 1994; Oremland, 1979; Ploug et al., 1997; Sieburth and Donaghay, 1993). These anoxic niches within floating algal aggregates can possibly serve as habitats for methanogenic archaea, thereby contributing to methane oversaturation in surface waters. While it is possible to evaluate a functional relationship with taxonomic sequence data alone, the correlation of the read number of Nodularia abundance and methanogenic Archaea suggests an environment favoring both groups of organisms simultaneously. The smooth algal mat structure of these sample types favors the development of a stable layered structure that would support the development of anoxic microniches.

The temporally community structure is in line with other observations (Mucko et al., 2020).

These algal mats may create complex metabolic networks where cyanobacterial production of dissolved organic carbon and extracellular polymeric substances provides essential substrates for methane production (Stuart et al., 2016). The physical structure of the algal mats enables the development of anoxic microsites, creating suitable conditions for methanogenic activity (Bižić et al., 2020b; Wilmeth et al., 2018). The stability of the mat community during all seasons suggests consistent methanogenic potential, with only minor variations in relative abundances indicating robust and well-established metabolic networks (Kurth et al., 2021). The interaction between these habitats creates important vertical coupling of processes (Baustian et al., n.d.; Griffiths et al., 2017). Seasonal mixing affects substrate availability across all zones, while temperature-dependent metabolic rates influence process intensities throughout the system (Tilstra et al., 2018). The most intensive methane-related processes likely occur in mat communities due to potentially more stable anoxic microsites, and among free-living communities through direct substrate utilization (Diaz et al., 2021). This could create a complex web of interactions where methane production potential may vary both spatially and temporally across these distinct but interconnected habitats. In support of the strong heterotrophic character of the algal mat samples is also the fact that the 18S V4 sequences were dominated by heterotrophs, and mostly metazoans, indicating that these communities support complex food webs with diverse metabolic interactions.

## Is there a relationship between season and methanogenic sequence/read number/mcrA abundance?

The seasonal variations affect the water column communities more than the algal mat communities, which maintain more stable year-round microbial abundances. This suggests that methane production might be more consistent in the algal mats but shows greater temporal variability in the water column, especially during periods of community transition (Bižić et al., 2020a; Peacock et al., 2023). The temporal community structure observed is in line with other observations from similar Baltic Sea environments (Mucko et al., 2020). The coupling between different functional groups also varies seasonally, particularly in the water column fractions, where changing environmental conditions influence the strength of metabolic interactions (Bižić et al., 2020a; Martinez-Cruz et al., 2015; Żygadłowska et al., 2023). The water samples show larger variations, which in the context of shallow inshore areas of the Baltic Sea may be related to potential mixing events including wind, water temperature changes, and algal blooms. The July-August period shows more consistent patterns across 0.22μm and 10μm samples, which may indicate mixing event. This is the period when algal blooms occur; during this time, deep mixing events can act as a source of nutrients that fuel these blooms (Martinez-Cruz et al., 2015).

## Is there evidence for a methane cycle close to the water surface characterized by methanogenesis and methane oxidation?

The 0.22μm water column fraction represents a more diverse community of smaller organisms, including free-living bacteria and archaea (Hahn, 2004; Li et al., 2021). Specifically, this fraction was dominated by Actinobacteriota, Bacteroidota, Cyanobacteria, Proteobacteria, and Verrucomicrobiota. Proteobacteria dominated from June to October, showing particularly high and stable proportions in habitat A, while being more variable in the other habitats. Actinobacteria and Bacteroidetes displayed distinct seasonal variation with peak abundances during the late summer months of August and September. In the filtered water samples, especially those from the 0.22 μm fraction, a more diverse array of methanogens was detected, including members of the Methanobacteriales order such as Methanobacterium and Methanothermobacter, which are capable of both hydrogenotrophic and methylotrophic methanogenesis and exhibit more diverse metabolic processes with a higher proportion of heterotrophic activity and potential for methyl compound metabolism (Leu et al., 2022). Additionally, ANME-3 archaea were present during multiple sampling periods in habitats A, C, and D, and anaerobic, sulfate-reducing bacteria belonging to the phylum Desulfobacterota, specifically Desulfobulbus clades, were detected with relative abundance less than 7.0%, suggesting potential anaerobic methane oxidation processes. The community composition shows distinct seasonal dynamics, with metabolic networks shifting in response to changing substrate availability. These patterns suggest that both filter size and habitat type play important roles in shaping bacterial community composition, with some phyla showing strong preferences for free-living conditions as indicated by their abundance in the 0.22 μm fraction, while others are more associated with particles larger than 10 μm, and generalists that maintain consistent distributions in all size fractions, seasons, and habitats.

In the algal mat habitat, the dominant presence of cyanobacteria indicates photosynthetic carbon fixation potential throughout the year (Wilmeth et al., 2018).

The shallow inshore habitats provided favorable environmental conditions for the growth and proliferation of various plants and benthic organisms during the summer and fall seasons, which thrive in the warmer temperatures and abundant nutrition sources available in these habitats. The presence of algal mats on the water’s surface created anoxic microenvironments that supported the growth of archaea. The most substantial methane fluxes were recorded in habitat A, particularly during the warmest sampling months of July and August (supplementary S2).

In the data set from the V3V4 amplicon sequence analysis using bacteria primer, we have observed the presence of three strains of cyanobacteria that Zou et al., (2024) detected in their study and associated with methane production. These are Nodularia PCC 9350, Synechocystis sp. PCC 6803, and Trichodesmium and they were consistently detected across various sample types, periods, and habitats. This consistency suggests that these strains are widespread and possibly play an important role in methane production across different environments. It’s important to note that while our study detected mcrA gene expression, indicating the presence of methane-producing microorganisms, this does not directly prove that these cyanobacterial strains are responsible for the methane production. The co-occurrence of these cyanobacterial strains with mcrA expression could also suggest a potential indirect role in methane production, possibly through the creation of favorable conditions or provision of substrates for methanogenic archaea, rather than direct methane production by the cyanobacteria themselves

Nodularia PCC-9350 showed the highest relative abundance among the three strains compared, particularly in 10 µm filtered water samples, indicating its presence in the free water column. This organism falls within the Cyanobacteria phylum and has been documented to consume naturally occurring phosphonates, displaying a particular affinity for methylphosphonate. This consumption has been linked to the generation of methane within the Baltic Sea, as outlined in a study conducted by Teikari et al., (2018).

This co-presence of Nodularia PCC-9350 raises questions about the nature of their relationship. While the data demonstrates their simultaneous existence in these environments, the exact nature of their interaction, whether symbiotic or independent, remains unclear and warrants further investigation. Multiple possibilities could explain this co-occurrence. One hypothesis is that a symbiotic relationship may be present, where Nodularia creates favorable microenvironments for methanogens. As a photosynthetic organism, Nodularia could potentially generate oxygen-depleted microzones within algal aggregates, providing suitable anaerobic habitats for methanogens. Additionally, Nodularia may release organic compounds that serve as substrates for methanogenic activity.

Alternatively, the co-presence of Nodularia and methanogens might be independent, simply reflecting their ability to thrive under similar environmental conditions in the shallow inshore habitats studied. Another consideration is Nodularia PCC 9350’s ability to consume methylphosphonate, which has been linked to methane generation in previous studies. This suggests a potential indirect relationship, where Nodularia’s metabolic activities influence methane production pathways, even in the absence of direct symbiosis.

To definitively establish the nature of this relationship, further targeted research is necessary. Future studies could employ co-culture experiments, fine-scale spatial distribution analyses, or metagenomic approaches to elucidate potential metabolic interactions. Examining the temporal dynamics of Nodularia and methanogen abundances could also provide insights into whether their populations follow similar patterns, potentially indicating a more direct relationship.

Understanding the precise nature of this co-occurrence is crucial for accurately interpreting methane dynamics in these aquatic ecosystems and could have significant implications for modeling methane emissions from shallow inshore habitats.

In summary, in Baltic Sea shallow waters, Nodularia and methanogens can interact through processes like anoxic zone creation, nutrient cycling, and seasonal dynamics, but maintain independent existence. However, within algal mats, their relationship becomes more intimate due to the algal mat’s layered structure, which creates distinct oxygen gradients and enables direct metabolic exchange. This difference suggests that filtered water samples would show loose associations, while mat samples would reveal stronger functional linkages between these organisms.

This formulation acknowledges the observed co-occurrence, presents multiple possibilities for the nature of the relationship, and suggests future research directions to clarify this aspect of the study. It also ties the discussion back to the broader implications for understanding methane dynamics in these ecosystems.

We also detected other already known methane producing cyanobacteria by Bižić et al., (2020b) such as Leptolyngbya, Prochlorococcus, Synechococcus, Trichodesmium, Anabaena, Nodularia, Scytonema, and Phormidium. Furthermore Synechocystis and Cyanobium were also identified, which belongs to picoplankton cyanobacteria together with Synechococcus and Prochlorococcus, and which have been identified participating in oxic methane production (Fazi et al., 2021; Tenorio and Farías, 2024). The presence of freshwater cyanobacteria such as Synechocystis sp. PCC 6803, Anabaena, and Scytonema across all habitats suggests some adaptation to brackish conditions. This suggests that methane production in Baltic Sea environments must consider both surface water production by cyanobacteria in addition to methane produced by archaea both in surface water floating algal mats and in sediment.

Furthermore, our analysis revealed a notably low presence, accounting for less than 0.5% relative abundance of Vibrio strains in all examined sample types and habitats. It is pertinent to highlight that Vibrio groups possess the potential for methane production. However, their occurrence in the Baltic Sea appears to be relatively limited, as previously reported by Eiler et al., (2006).

The investigation of the presence of iron, nitrate and sulfate reducers showed that for the most part these occur at low levels of relative abundance, and all three types co-occurred in habitat A in July and September in filtered water samples.

The presence of a 16S community indicative of iron and nitrate reducers co-occurring with sulfate reducers is possible because microbial communities, including bacteria involved in various biogeochemical processes, such as methane production, which can co-occur with other metabolic processes like iron, manganese, and sulfate reduction in oxygenated surface waters. This coexistence suggests that complex microbial interactions are at play in these environments (Tenorio and Farías, 2024), which fit well with the characteristics of habitat A.

The notably higher values of nitrate reducers in algal mat samples may indicate fluctuating oxygen conditions (Sørensen, 1982), and possibly vertically stratified redox conditions.

In our results we had detected Desulfobulbus in both 10 µm and 0.22 µm filtered water samples. This genus is frequently associated with anaerobic methanotrophic archaea, ANME, (Bhattarai et al., 2017). The finding of Desulfobulbus suggests the possibility of a dependent anaerobic oxidation of methane by ANME-3 archaea. The results show the presence of ANME-3 archaea during multiple sampling periods, and in habitats A, C, and D. The presence of ANME-3 archaea in particularly in filtered water samples, habitat C, is quite interesting, since this group of archaea exists in coastal marine sediment and is involved in the process of anaerobic methane oxidation (Bhattarai et al., 2017). This process converts methane gas into different compounds without the need for oxygen, playing a crucial role in reducing methane emissions. As noted by Bhattarai et al., (2017), presence of ANME-3 in shallow coastal areas has not been commonly reported, and more commonly found at submarine mud volcanoes and to some extent cold seeps (Vigneron et al., 2013).

This intriguing result indicates that the anoxic microenvironments in the surface water may provide suitable conditions for the anaerobic oxidation of methane, although an alternative explanation cannot be ruled out. It is important to note that our sampling region encompasses shallow inshore areas with water depths ranging from 0.5 to 2.0 meters. It is possible that boat movements and sample collection activities could release these microorganisms from the sediment in the shallowest areas (A and B), although this remains speculative, as ANME-3 has generally been detected a few centimeters below the sediment surface (Vigneron et al., 2013), and no sediment samples were collected during our study. Also, for habitat C the water depths were generally on the higher end of the range and thus less likely to have caused disturbances of the sediment. Bhattarai et al., (2017) identified ANME-3 archaea as the main drivers of anaerobic methane oxidation in this sediment environment. These archaea collaborate with sulfate-reducing bacteria to convert methane into more stable compounds. This research sheds light on the complex interactions between microorganisms and the processes that contribute to methane cycling in marine sediments, offering insights into the carbon and methane dynamics in coastal ecosystems.

Notably though, Desulfobulbus were found also when there was no presence of ANME-3, in the filtered water samples. This suggests that it may thrive in localized low-oxygen or anaerobic environments, and the presence in 0.22 µm filtered water samples that it may exist as free-living planktonic bacteria.

These results comparing ANME-3 and Desulfobulbus suggests that the latter may have a broader adaptability from an ecological perspective, being able to survive in various oxygenic conditions across different habitats (Seidel et al., 2023). *Desulfobulbus* is a sulfate-reducing bacterium adapted to anoxic environments, with its distribution and diversity influenced by temperature and sediment depth, and potentially engaging in syntrophic relationships with archaea under certain conditions (Seidel et al., 2023). ANME-3’s presence is more limited and potentially be limited by environmental disturbances as well.

Recent research has identified Methanolobus as a key participant in anaerobic dimethylsulphide degradation in brackish Baltic Sea environments (Tsola et al., 2024). In our study, Methanolobus was detected in habitat A and B, but only in 10 µm filtered water samples, appearing in September and August respectively. Notably, the co-occurrence of Methanolobus, ANME-3, and Desulfobulbus was observed only in habitat A during September, which could strengthen the likelihood of a sediment signal in our samples for this case.

Analysis of the methanogenic archaea communities across different habitats and sampling periods reveals patterns that allow for speculation on the dominant methane production pathways in these environments. Examination of the taxonomic classification and known metabolic capabilities of the detected methanogens provides insights into potential active methanogenesis routes in the samples and is instrumental in discerning the specific methanogenic pathways employed for methane production. Our findings provide valuable insights, indicating the presence of both hydrogenotrophic and methylotrophic methanogenic groups within oxic water environments.

In a previous study, Methanomicrobiales were identified as a hydrogenotrophic methanogenic community residing within smooth algal mats (Wong et al., 2017). Our analysis reveals the presence of this community predominantly in algal mat samples collected from habitat A, up to 100% of methanogenic archaea. However, the genus Methanocorpusculum belongs to this order, and appears in relatively lower abundance in samples from habitats B and D during the sampling period of July and August for algal mats. Interestingly, Methanomicrobiales were also detected in filter samples with a pore size of 0.22 µm from the same habitats and sampling periods. Since these organisms are known hydrogenotrophic methanogens, this confirms that hydrogenotrophic methanogenic pathway occurs both in algal mat and in phytoplankton, and suggesting that the hydrogen-dependent pathway of methanogenesis may be significant in these surface-associated microbial communities (Browne et al., 2017; Joshi et al., 2018). The presence of these hydrogenotrophs in algal mats could indicate localized production of hydrogen, possibly from fermentative processes occurring within anoxic micro niches of the mats (Volmer et al., 2023). Conversely, in the filtered water samples, especially those from the 0.22 μm fraction, a more diverse array of methanogens was detected, including members of the Methanobacteriales order such as Methanobacterium and Methanothermobacter. These groups are capable of both hydrogenotrophic and methylotrophic methanogenesis, suggesting a potential shift in substrate utilization in the free-living microbial communities compared to those in the algal mats (Kurth et al., 2020; Lv et al., 2022). The substantial presence of Methanosarcina suggests multiple pathways are active simultaneously, while the consistent detection of Methanobacterium indicates ongoing hydrogenotrophic methanogenesis (Milici et al., 2016; Ziganshin et al., 2016). The periodic appearance of Methanosaeta indicate aceticlastic methanogenesis activity during specific timepoints (Chen et al., 2017; Mori et al., 2012).

Notably, a temporal variation in the methanogen community composition was observed. During warmer sampling periods (July and August), there was a notable increase in the relative abundance of Methanosarcina in habitats A, B, and D, particularly in the 10 μm filtered samples. Methanosarcina is metabolically versatile, capable of all three main methanogenesis pathways: hydrogenotrophic, methylotrophic, and acetoclastic (Ferry, 2011). This shift towards Methanosarcina dominance during warmer periods might indicate a diversification of available substrates, possibly due to increased organic matter decomposition at higher temperatures (Reichlen et al., 2012). The filter 10 µm samples display less diverse communities but exhibit pronounced temporal shifts, with later periods demonstrating strong dominance by either Methanobacterium or Methanosarcina, suggesting shifts between primarily hydrogenotrophic methanogenesis and potentially more diverse pathway utilization respectively (Milici et al., 2016; Ziganshin et al., 2016). The presence and abundance of dominant genera provide interesting insights into community structure, as Methanobacterium, Methanocella, and Methanosarcina emerge as key players, but their relative abundance fluctuate considerably across samples. This could reflect their different ecological niches and responses to changing environmental conditions (Liu et al., 2024). Some samples show near-complete dominance by a single genus, suggesting conditions that strongly favor particular methanogenic groups (Liu et al., 2024).

The reported methylotrophic methanogenic groups; Methanosarciniales (Wong et al., 2017) and Methanomassiliicoccales are present only in the algal mat samples, in our data, while Methanobacteriales is present in all sample types. More specifically, for Methanomassiliicoccales we detected Candidatus Methanomethylophilus in habitat A and B, which suggests the potential importance of methylated compound metabolism in these environments, which is also interesting given the co-occurrence Nodularia PCC 9350, which is known to produce methylated compounds that could serve as substrates for these methanogens. The algal mat samples from July and August indicate a dominance of hydrogenotrophic methanogenesis, possibly driven by the high abundance of Methanobacterium (Quéméneur et al., 2021; Sonne-Hansen and Kiær Ahring, 1997). This pattern shifts in September, where we observe increased community diversity and a notable rise in Methanosarcina abundance, suggesting a broader range of active methanogenic pathways including aceticlastic and methylotrophic methanogenesis alongside hydrogenotrophic pathways(Aschenbach et al., 2013; Torres-Alvarado et al., 2013; Wong et al., 2017). Methanobacterium, which belong to order Methanobacteriales, were detected in algal mat samples in habitat D (Figure 5-a), and in filter 0.22 µm in all habitats (Figure 5-c). Methanothermobacter, which also belong to order Methanobacteriales were detected in filter pore size 0.22 µm and 10.0 µm (Figure 5-b, c). This indicates that methylotrophic methanogenic pathway occurs both in algal mat and in phytoplankton.

However, in habitat D, Methanosarcina and Methanobacterium, which belong to order Methanobacteriales, dominated the algal mat samples. Specifically, Methanosarcina was found in both habitat D and B, but was more abundant in habitat D, making up 67.2% relative abundance of methanogenic archaea there, compared to 1.3% relative abundance in habitat B. Considering the hard unvegetated sediment in habitat D beside the high relative abundance of Methanosarcina, we can conclude the active methane production in algal mats in habitat D. This finding confirms the qPCR results with the highest level of methanogenic mcrA gene expression in algal mat samples collected from habitat D.

All algal mat samples collected in our study are smooth mats according to the categorization in (Wong et al., 2017).The study by Wong et al., (2017) revealed noteworthy variations in metabolic activities between smooth and pustular microbial mats. Smooth mats exhibited significantly elevated rates of both oxygen production/consumption and sulfate reduction in comparison to their pustular counterparts, as documented in the study by Wong et al., (2017). Furthermore, in the same study, methane production reached its peak in the oxygen-rich layers, with smooth mats displaying methane production levels up to seven times greater than pustular mats. This phenomenon suggests the existence of surface anoxic niches within smooth mats, where anaerobic methanogens thrived despite the prevailing oxygen-rich conditions. These findings underscore the intricate nature of microbial interactions within these ecosystems, especially among diverse archaeal groups, and provide valuable insights into the finely tuned spatial dynamics at play.

These observations indicate a potential spatial separation of methanogenic pathways, with hydrogenotrophic methanogenesis possibly dominating in algal mats, while a more diverse set of pathways, including methylotrophic and potentially acetoclastic methanogenesis, may be active in the water column. The temporal shifts in community composition further suggest that the relative importance of these pathways may vary seasonally, with a trend towards more metabolically versatile methanogens during warmer periods. The frequent dominance of Methanosarcina suggests metabolic flexibility plays a crucial role in these environments. However, it’s important to note that while these abundance patterns suggest potential pathway activity, definitive confirmation would require additional molecular and biochemical analyses such as substrate measurements, gene expression studies, or isotope labeling experiments. These observations align with current understanding of methanogenic community dynamics in anaerobic environments, where substrate availability and environmental conditions often drive shifts in dominant pathways and community composition. The temporal variations observed suggest dynamic responses to changing environmental conditions or substrate availability patterns across the sampling periods. Habitat specificity is another notable pattern, with different sampling locations showing distinct methanogenic communities. This spatial variation indicates that local environmental conditions play a crucial role in shaping community composition (Jing et al., 2016). The algal mat samples, in particular, show unique community profiles compared to the filtered samples, suggesting that these habitats provide specific niches for certain methanogenic groups (King, 1988).

Another consideration is that the methanogenic archaeal communities show distinct temporal patterns that may be driven by both physical mixing events and biological processes. During the summer months (July-August), when water temperatures were stable at 15-17°C and algal blooms typically occur in the Baltic Sea, the community composition suggests active processing of fresh organic matter (Thureborn et al., 2013; Wäge et al., 2020). This is indicated by the presence of both aceticlastic (Methanosaeta, Methanosarcina) and hydrogenotrophic (Methanobacterium) methanogens, along with elevated methane concentrations in late July. Methanobacterium shows particularly strong representation during the July-August period, indicating potential environmental conditions that favor their growth during these months (Fu et al., 2023).

The most pronounced community shifts occurred in September, coinciding with both post-bloom conditions and the first significant temperature drop. These changes were accompanied by peak methane concentrations, suggesting intense decomposition activity (Lundevall-Zara et al., 2021; Thureborn et al., 2013). The parallel shifts in community composition across different size fractions (0.22 and 10 µm filters) and algal mat samples indicate system-wide changes affecting both particle-attached and free-living communities. While moderate wind conditions (3-6 m/s) could contribute to water column mixing in these shallow (0.5-2.0 m) areas, the observed succession in methanogenic communities may be equally influenced by changing substrate availability as algal biomass decomposes, progressing from initial acetate availability to more complex organic matter degradation pathways (Thureborn et al., 2013).

These speculations provide a framework for understanding the complex dynamics of methane production in these shallow inshore habitats. However, further research, such as stable isotope probing or metatranscriptomic analyses, would be necessary to conclusively determine the active methanogenic pathways and their relative contributions to overall methane production in these ecosystems. These patterns collectively suggest a complex interplay between temporal, spatial, and methodological factors in determining the structure of methanogenic archaeal communities in these environments (Jing et al., 2016). The variations observed likely reflect both the ecological adaptations of different methanogenic genera and their responses to changing environmental conditions (Jing et al., 2016; White, 2023).

The study employed PERMANOVA to assess significant differences in multivariate data based on group assignments. For the 16S V4V5 Archaea and 16S V3V4 Bacteria datasets, the analysis revealed moderate differences based on sampling period in the archaeal community for the 10 μm and 0.22 μm filter datasets. This suggests that the sampling period has a more substantial impact on Archaeal and Bacterial composition in filter samples. This effect may potentially be due to seasonal variations (e.g. temperature, nutrients, light) and hydrological changes (e.g. water mixing and flow), possibly to a larger degree than for algal mats. However, given that the results for the algal mat PERMANOVA have high P-values, the significance of the R^2^ values for these samples are inconclusive.

These findings align with NMDS analysis, which also indicated more pronounced differences between sampling periods for filter samples compared to algal mats. Furthermore, the PERMANOVA tests for the 10 μm and 0.22 μm filter samples demonstrated higher R-squared values than the test for algal mats.

By identifying various methane-producing microorganisms, such as Cyanobacteria and methanogenic Archaea, the research highlights the critical role these habitats play, especially in the presence of algal mats which foster both oxygen-rich and anoxic conditions conducive to methane production. Findings suggest that human activities affecting these coastal areas, coupled with natural processes, significantly contribute to atmospheric methane levels (Rocher-Ros et al., 2023). Therefore, it is essential to include aquatic methane emissions from these regions in the global methane budget to better understand and mitigate their impact on climate change.

The Delta CT values in Table 1 are lower for algal mat samples compared to filtered water samples, which would indicate higher mcrA gene expression of the free-living microbiota, compared to algal mats. This is a bit of a surprising result. However, the methane concentrations also vary even with similar Delta CT values, so there is not a clear correlation between methane concentration and mcrA gene expression levels. This may suggest that there may be multiple methanogenic pathways involved (Wang et al., 2022; Zhang et al., 2019), additional environmental factors (Wang et al., 2022) or microbial composition factors that are at play here. For example, acetoclastic methanogenesis pathway would be affected by acetate availability or presence of other acetate-utilizing microbes, since this may have an impact on the abundance of acetoclastic methanogens (Wang et al., 2022). This in turn may result in variations in Delta CT values. The taxonomic composition analysis includes the detection of both sulfate-reducing bacteria e.g. Desulfovibrio and Desulfobacter, syntrophic acetate-oxidizing bacteria, as well as Denitrying bacteria (e.g. Pseudomonas), which potentially could have affected results for acetoclastic methanogenesis pathway (Jiang et al., 2018; Mesquita et al., 2023; Wang et al., 2022). Furthermore, since we detected different genus of methane producing archaea and cyanobacteria in different sample types, we can conclude that it is not due to resuspension from sediment because if that was the case, we would expect to find the same genera combinations in multiple sample types.

**Table 1.** qPCR results compared to CH4 concentration for different sampling periods across all habitats.

| Sample name | Habitat | Station Name | Sampling date | Sampling Period | Flux (mg/m<sup>2</sup>/day) | Mean RT-qPCR 16S CT | Mean RT-qPCR mcrA CT | Delta CT |
| --- | --- | --- | --- | --- | --- | --- | --- | --- |
| Algal mat |  |  |  |  |  |  |  |  |
| 32-2-Algal | A | 2(2-E) | 2020-07-09 | Jul |  | 23.68 | 33.10 | 9.43 |
| 34-2-Algal | A | 2(2-G) | 2020-07-09 | Jul |  | 30.70 | 32.87 | 2.16 |
| 36-8-Algal | B | 8(1-A) | 2020-07-07 | Jul | 0.19 | 31.38 | 32.63 | 1.25 |
| 31-8-Algal | B | 8(1-C) | 2020-07-07 | Jul | 0.19 | 30.71 | 34.24 | 3.53 |
| 33-3-Algal | D | 3(4-M) | 2020-08-27 | Aug |  | 31.38 | 34.52 | 3.14 |
| 30-3-Algal | D | 3(4-N) | 2020-08-27 | Aug | 0.14 | 33.40 | 34.84 | 1.45 |
| Filter 0.22µm |  |  |  |  |  |  |  |  |
| 14-7-3-0.22 | A | 5(5-V) | 2020-07-08 | Jul |  | 13.47 | 33.78 | 20.30 |
| 21-7-3-0.22 | A | 7(5-V) | 2020-08-26 | Aug |  | 23.68 | 33.89 | 10.21 |
| 29-7-1-0.22 | A | 7(5-V) | 2020-08-26 | Aug |  | 18.82 | 33.73 | 14.90 |
| 17-2-1-0.22 | A | 2(2-F) | 2020-08-27 | Aug | 0.52 | 12.21 | 34.92 | 22.70 |
| 22-8-1-0.22 | B | 8(1-C) | 2020-08-27 | Aug | 0.07 | 17.64 | 33.35 | 15.71 |
| 43-8-5-0.22 | B | 8(1-C) | 2020-08-27 | Aug | 0.07 | 20.58 | 32.95 | 12.38 |
| 7-3-1-0.22 | D | 3(4-N) | 2020-07-08 | Jul | 0.25 | 17.84 | 33.84 | 15.99 |
| 12-6-1-0.22 | D | 6(5-U) | 2020-07-08 | Jul |  | 13.77 | 33.91 | 20.14 |
| 18-3-3-0.22 | D | 3(4-N) | 2020-08-27 | Aug | 0.14 | 18.92 | 33.83 | 14.90 |
| Filter 10µm |  |  |  |  |  |  |  |  |
| 1-1-1-0.10 | A | 1(3-I) | 2020-07-07 | Jul | 8.30 | 13.77 | 33.94 | 20.17 |
| 2-2-4-10 | A | 2(2-F) | 2020-07-08 | Jul | 1.34 | 14.69 | 34.92 | 20.23 |
| 38-2-6-10 | A | 2(2-F) | 2020-07-08 | Jul | 1.34 | 18.52 | 33.34 | 14.81 |
| 10-5-2-10 | A | 5(5-Q) | 2020-07-08 | Jul |  | 17.53 | 34.27 | 16.75 |
| 40-7-4-10 | A | 7(5-V) | 2020-07-08 | Jul |  | 19.63 | 34.30 | 14.67 |
| 26-5-4-10 | A | 5(5-Q) | 2020-08-26 | Aug |  | 21.54 | 33.92 | 12.38 |
| 23-1-2-10 | A | 1(3-I) | 2020-08-27 | Aug | 3.16 | 19.23 | 33.37 | 14.14 |
| 16-8-6-10 | B | 8(1-C) | 2020-08-27 | Aug | 0.07 | 19.79 | 33.39 | 13.60 |
| 42-8-2-10 | B | 8(1-C) | 2020-08-27 | Aug | 0.07 | 19.19 | 33.01 | 13.81 |
| 8-4-2-10 | C | 4(4-P) | 2020-07-08 | Jul | 0.01 | 18.61 | 33.33 | 14.71 |
| 25-4-2-10 | C | 4(4-P) | 2020-08-26 | Aug | 0.88 | 28.93 | 34.35 | 5.42 |
| 39-3-2-10 | D | 3(4-N) | 2020-07-08 | Jul | 0.25 | 17.06 | 34.08 | 17.02 |
| 11-6-4-10 | D | 6(5-U) | 2020-07-08 | Jul |  | 15.52 | 33.66 | 18.15 |
| 27-6-2-10 | D | 6(5-U) | 2020-08-26 | Aug |  | 17.87 | 34.69 | 16.82 |
| 24-3-4-10 | D | 3(4-N) | 2020-08-27 | Aug | 0.14 | 27.93 | 34.59 | 6.66 |

**Table 2.** PERMANOVA results for V4V5 primer samples for archaeal community.

| Primer + sample type | Group variable | Degrees of freedom | R<sup>2</sup> | F-statistic | P-value |
| --- | --- | --- | --- | --- | --- |
| V4V5 Archaea algal mats | Sampling period | 2 | 0,12548 | 0,7174 | 0,849 |
| V4V5 Archaea filter 0.22 $\mu\text{m}$ | Sampling period | 4 | 0,33653 | 2,7897 | 0,001 |
| V4V5 Archaea filter 10 $\mu\text{m}$ | Sampling period | 4 | 0,30059 | 2,0056 | 0,038 |
| V3V4 Bacteria algal mats | Sampling period | 2 | 0,12697 | 0,7999 | 0,698 |
| V3V4 Bacteria filter 0.22 $\mu\text{m}$ | Sampling period | 4 | 0,21025 | 1,5974 | 0,034 |
| V3V4 Bacteria filter 10 $\mu\text{m}$ | Sampling period | 4 | 0,23059 | 1,948 | 0,018 |

## Conclusions

The long-held belief that methane production exclusively occurs in oxygen-deprived sediment environments through the actions of archaea is undergoing a significant revision. Multiple studies, including ours, have presented compelling evidence supporting methane production in surface oxygen-rich waters. Our qPCR analysis provides clear and significant confirmation, highlighting the highest levels of methanogenic mcrA gene expression within algal mat samples. Interestingly, even our filtered water samples exhibit active methanogenic mcrA gene expression. This discovery carries crucial implications, particularly concerning the roles of eutrophication and global warming in amplifying methane production within these shallow habitats. The combination of global warming and eutrophication leads to greater vegetation growth, including algal mats and blooms, in the warmest months. This, in turn, results in elevated methane emissions from shallow inshore regions during the warmest periods of the year. The findings of this study offer insights into the underlying metabolic processes in these areas. In conclusion, it’s likely that methane production in these habitats is a combination of both sediment and water column processes, with potential contributions from both the photic zone and sediment-based sources. The relative importance of each source may vary depending on the specific characteristics of each habitat, including water depth, vegetation, algal mats presence and thickness, and sediment organic content. Further targeted studies would be necessary to quantify the exact contributions from each source in these specific habitats.

## Highlights and key points

Archaeal 16S and mcrA abundant in surface water samples of vegetated organic-rich habitats

Floating algal mats also contain methanogenic archaea, but not at higher levels than corresponding water samples.

The bacterial community contains abundant sequences of anaerobic microorganisms, the most notable sulfate-reducing SRB.

Organisms that are associated with the anaerobic oxidation of methane were also found indicating that organisms carrying out a complete anaerobic methane cycle are present in oxic surface environments.

Although supportive of active methanogenesis in photic, fully oxygenated surface waters, the absence of a good correlation between methane levels and sea-to-air methane flux indicates that methanogenesis in these surfaces likely does not have a dominating influence.

Instead, the abundance of ANME and Desulfobulbus in the 10 µm particle fraction can be interpreted as an important contribution of sediment suspension, possibly even further exacerbated by methane bubbling, for the surface water microbial community.

The distinct combinations of methanogenic archaeal and cyanobacterial genera found across different sample types suggest that their presence is not due to sediment resuspension, as such resuspension would have resulted in consistent genera combinations across multiple sample types.

## Data availability

The environmental data used in this study can be found at NCBI website (https://www.ncbi.nlm.nih.gov/) through BioProject database and accession number PRJNA1070012. https://dataview.ncbi.nlm.nih.gov/object/PRJNA1070012?reviewer=f2sfn0t2hnf7fpfa3div5cd251

## Acknowledgements

I am deeply grateful to my supervisor Volker Brüchert from Department of Geological Sciences for all the support and valuable insights into this work. Special thanks also go to Elias Broman at DEEP, Stockholm university, for methodological guidance, as well as the staff of Askö Laboratory for their assistance during the sampling efforts, and Francisco Nacimiento for providing access to the DEEP lab at Stockholm’s University.

## Notes

### Competing Interest Statement

The authors have declared no competing interest.

