## Supplemental material for "Methane-producing microorganisms are widespread in surface waters and floating algal mats of inshore Baltic Sea habitats"

Maysoon Lundevall Zara, Tibble Gymnasium Campus Täby

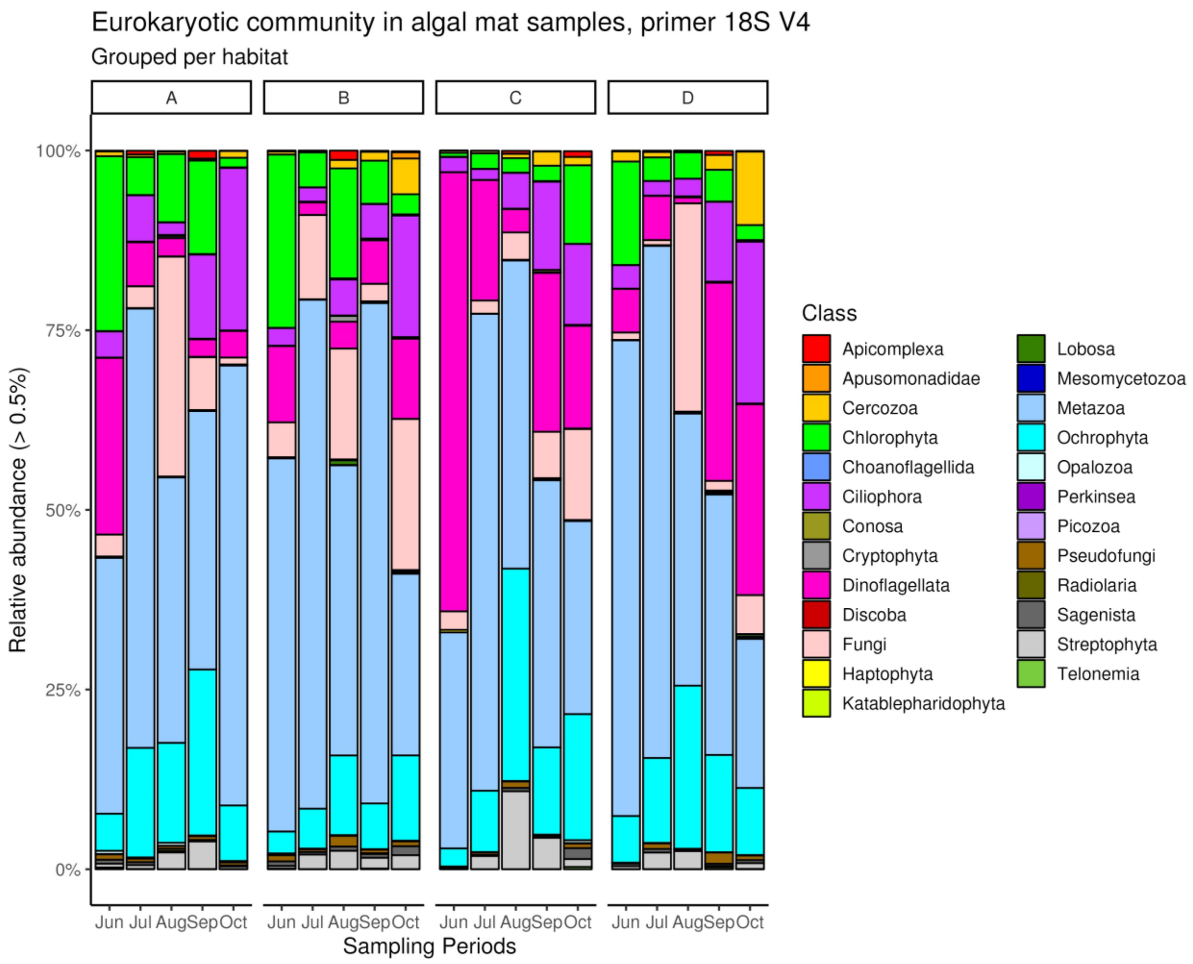


Figure S1: Eukaryotic community at class level grouped by habitat and sampling period in algal mats samples with 18S V4 primers.

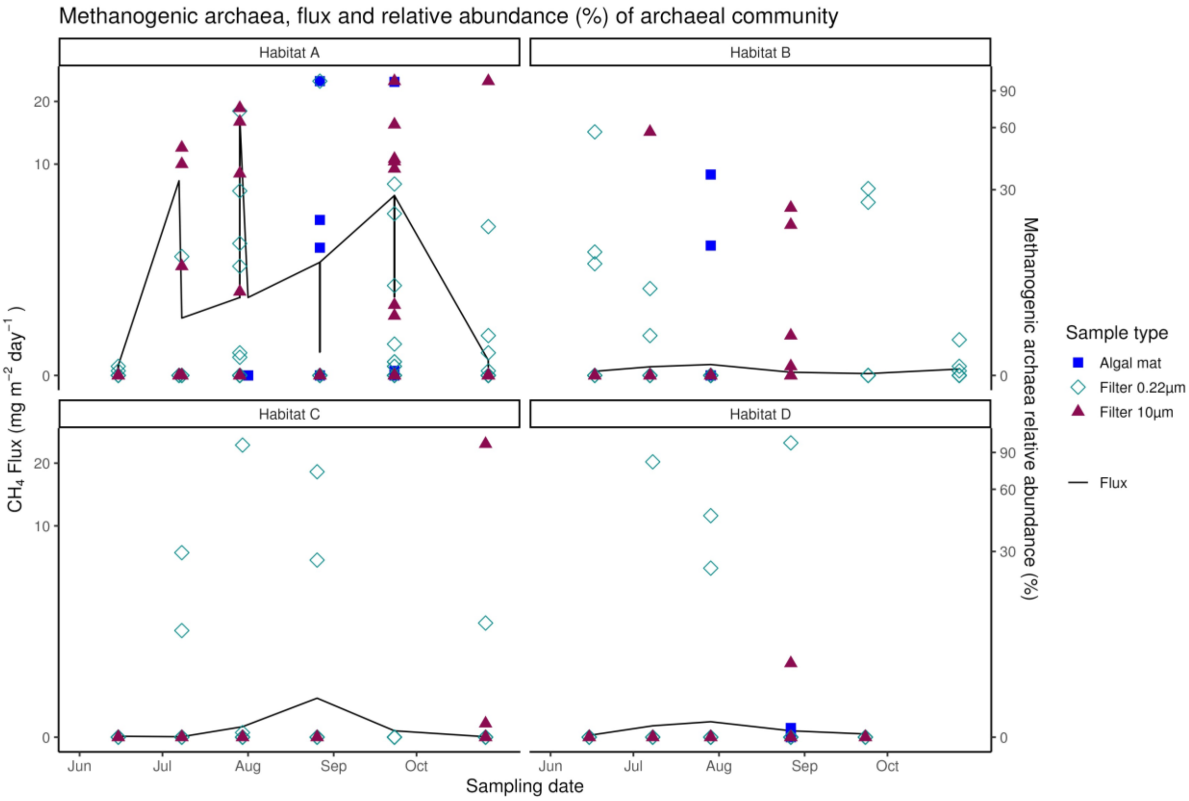


Figure S2: Methanogenic archaea, flux, and relative abundance (%) of archaeal community in all sampling types

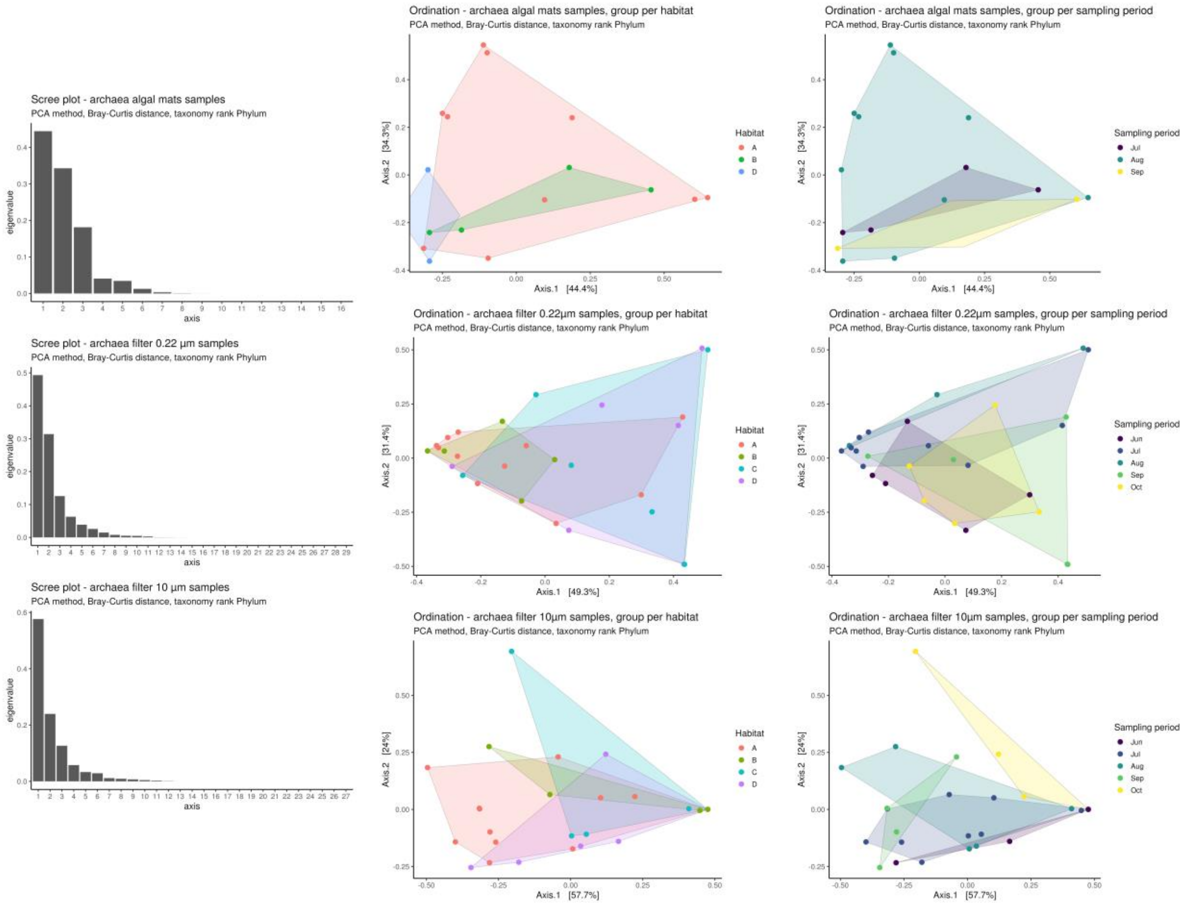

Figure S3: PCA analysis for different habitat and sampling periods for algal mat samples, filter 0.22µm samples and filter 10µm samples for Archaea primer.

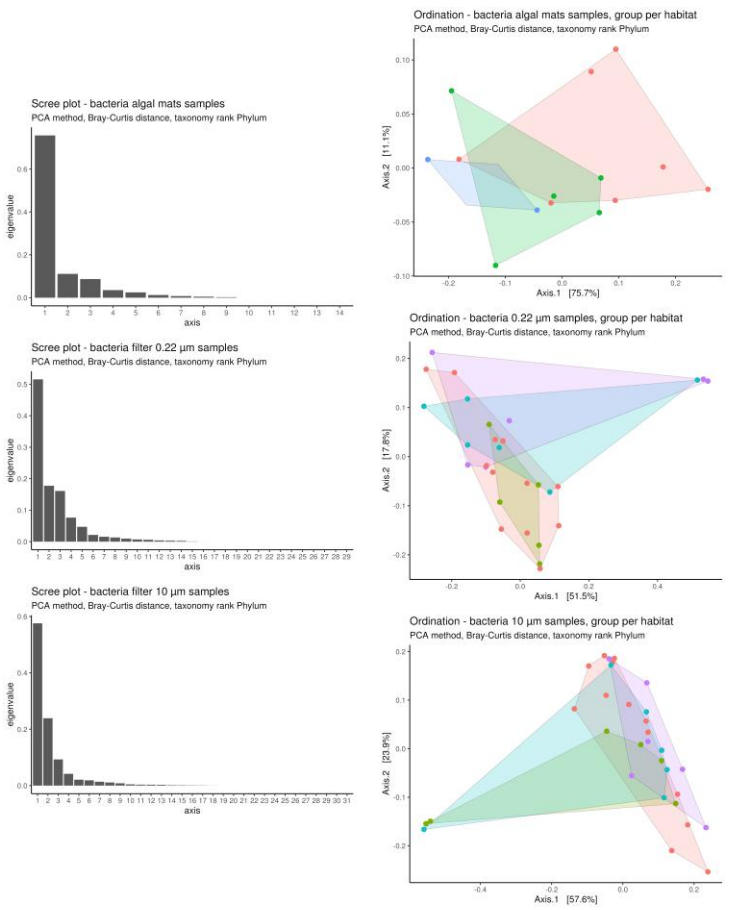


Figure S4: PCA analysis for different habitat and sampling periods for algal mat samples, filter 0.22µm samples and filter 10µm samples for Bacteria primer.

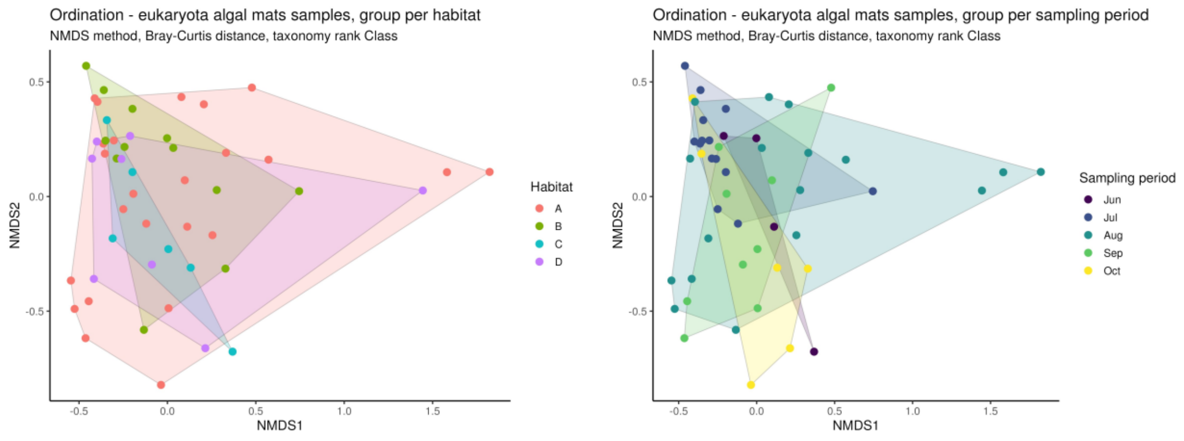


Figure S5: NMDS analysis for different habitat and sampling periods for algal mat samples for Eukaryote.

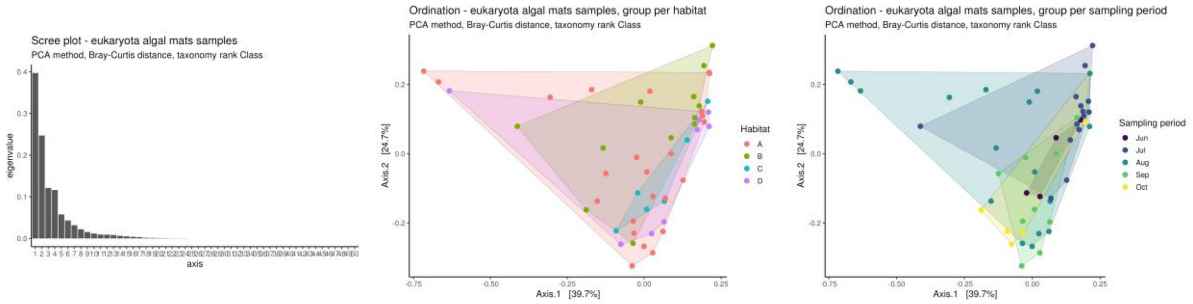


Figure 14: PCA analysis for different habitat and sampling periods for algal mat samples for Eukaryote.

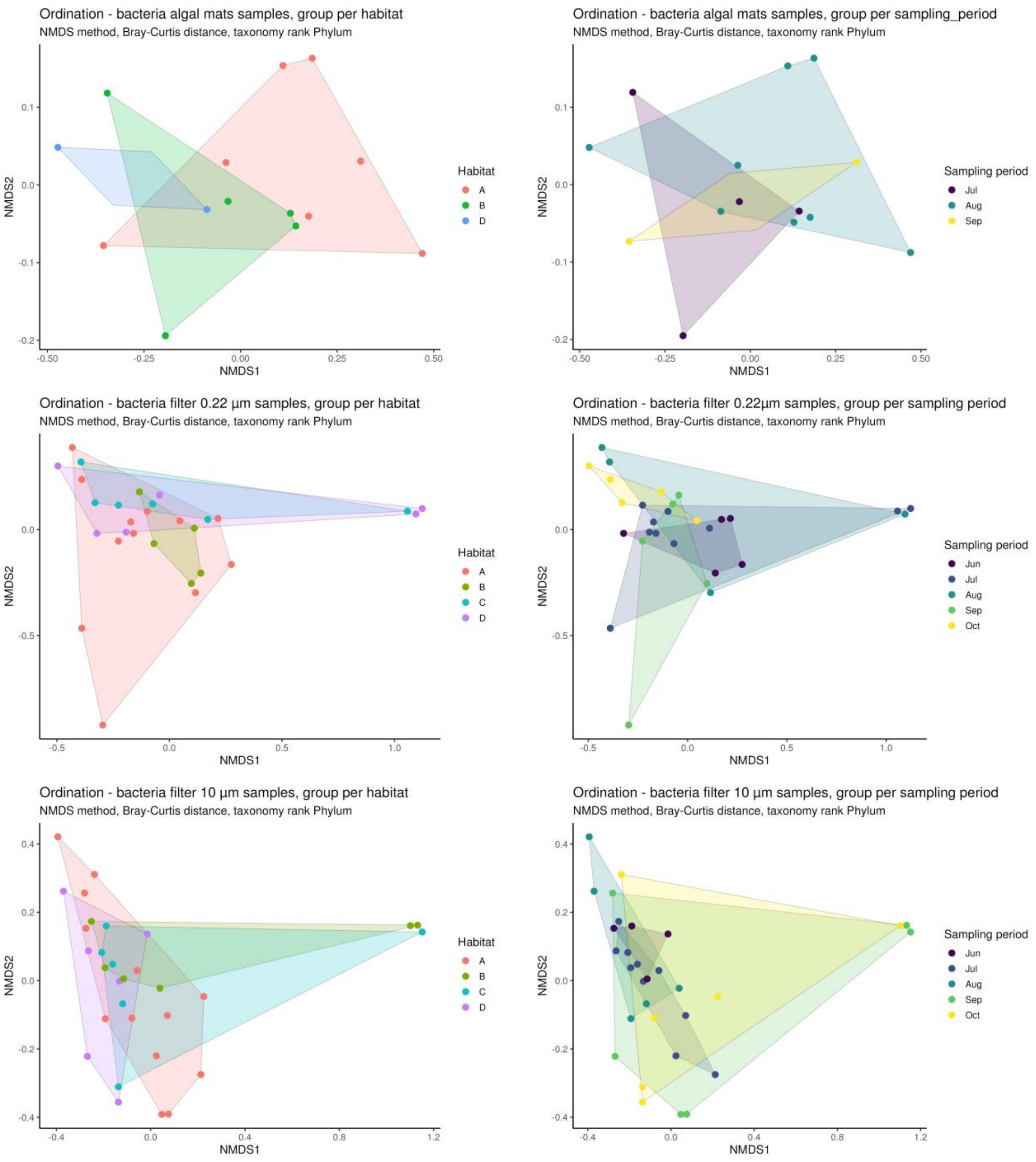


*Figure 1 NMDS analysis for different habitat and sampling periods for algal mat samples, filter 0.22µm samples and filter 10µm samples for Bacteria primer.*

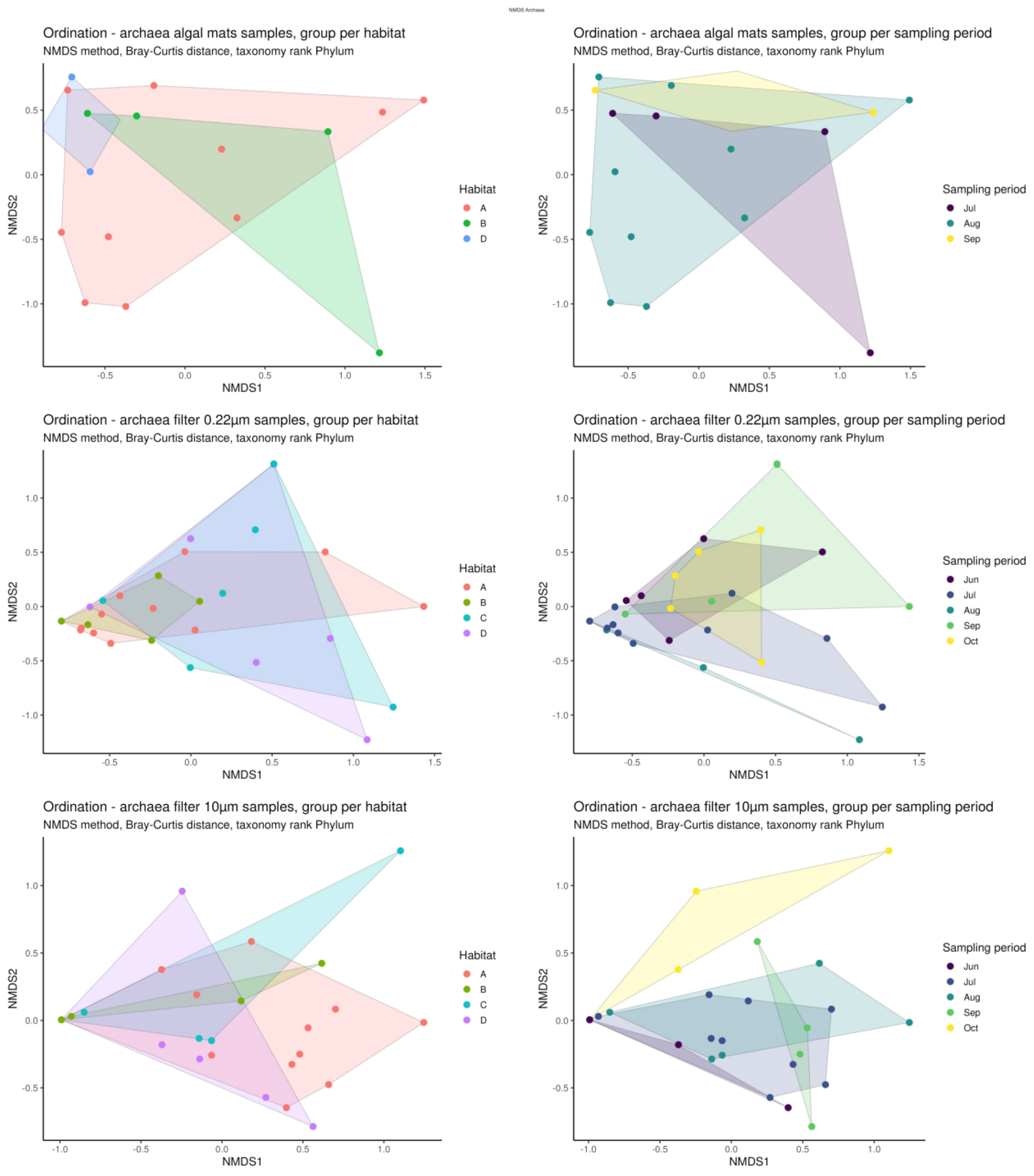


*Figure 2 NMDS analysis for different habitat and sampling periods for algal mat samples, filter 0.22µm samples and filter 10µm samples for Archaea primer.*
